# Down syndrome fibroblasts exhibit diminished autophagic clearance and endosomal dysfunction after serum starvation

**DOI:** 10.1101/436782

**Authors:** Stefanos Aivazidis, Abhilasha Jain, Colin C. Anderson, David J. Orlicky, Abhishek K. Rauniyar, Kristofer S. Fritz, Peter S. Harris, David Siegel, Kenneth N. Maclean, James R. Roede

**Affiliations:** Department of Pharmaceutical Sciences, Skaggs School of Pharmacy and Pharmaceutical Sciences, University of Colorado, Aurora, CO, USA; Department of Pathology, University of Colorado School of Medicine, Aurora, CO, USA; Department of Pediatrics, University of Colorado School of Medicine, Aurora, CO, USA; The Linda Crnic Institute for Down Syndrome, University of Colorado, Aurora, CO, USA

**Keywords:** Down syndrome, autophagy, aneuploidy, proteostasis network, LC3-II, p62, serum starvation, Rab5, Rab11

## Abstract

Down syndrome (DS) is a genetic disorder caused by trisomy of chromosome 21 (Tri21). This unbalanced karyotype has the ability to produce proteotoxic stress and dysfunction of the proteostasis network (PN), which are mechanistically associated with several comorbidities found in the DS phenotype. Autophagy is the cellular process responsible for bulk protein degradation and its impairment could negatively impact protein quality control. Based on our previous observations of PN disruption in DS, we investigated possible dysfunction of the autophagic machinery in human DS fibroblasts. Both euploid (CTL) and DS fibroblasts induced autophagy successfully through serum starvation (SS), as evidenced by increased LC3-II abundance in CTL and DS. However, DS cells displayed evidence of autophagolysosome (AL) accumulation and impaired clearance of autophagosome cargo, e.g. accumulation of p62 and NBR1. Similar observations were also present in DS cells from multiple differentiation stages, implicating impeded autophagic degradation as a possible early pathologic mechanism in DS. Lysosomal pH and cathepsin B proteolytic activity were found to not differ in CTL and DS fibroblasts after SS, indicating that lysosomal dysfunction did not appear to contribute to unsuccessful autophagic clearance. Based on these results, we hypothesized that possible interference of the endosomal system with autophagy results in autophagosome fusion with endosomal vesicles and negatively impacts degradation. Consistent with this hypothesis, we observed increased abundance of the recycling endosome marker, Rab11, after SS in DS fibroblasts. Further, colocalization of autophagosome markers with resident proteins of early endosomes, late endosomes and recycling endosomes (Rab11) further support our hypothesis. In summary, our work is consistent with impairment of autophagic flux and general PN dysfunction as candidate mechanisms for pathology in DS.

## Introduction

Down syndrome (DS) is a genetic condition originating from the presence of a third copy of chromosome 21 (Tri21) (*1*) and is the only known trisomy that does not result in early life lethality (*2*). DS is characterized by a variable phenotype with several comorbidities, including Alzheimer’s disease (AD) (*1*), cognitive disabilities (*3*), diabetes (*4*), and leukemia (*5–7*). Previous research from our laboratory and others has revealed the presence of a dysfunctional proteostasis network (PN) in DS cell models, characterized by several markers of disrupted proteostasis (*8–12*). More specifically, DS cells exhibit basal endoplasmic reticulum (ER) stress, increased abundance of polyubiquitinated proteins and decreased proteasomal degradation. These are critical observations as the PN serves as guardian of the proteome by preserving the integrity of protein biogenesis, folding and degradation (*13*). Importantly, PN dysfunction can act as a mechanism to promote pathology, and many comorbidities appearing in the DS phenotype are mechanistically associated with impaired proteostasis (*14–21*). Since PN dysfunction in DS has yet to be fully investigated, we present data here on the autophagic process in DS.

The mammalian cell displays three types of autophagy: 1) Microautophagy, in which the lysosome changes conformation of its membrane to create invaginations that capture cytosolic material (reviewed here (*22*)); 2) Chaperone-mediated autophagy, in which the constitutively expressed isoform of the HSP70 chaperone family (Hsc70) binds and shuttles individual misfolded peptides to the lysosome, where they undergo proteolytic degradation. (reviewed here(*23*)); 3) Macroautophagy (hereafter referred to as autophagy), which is responsible for bulk protein degradation and elimination of dysfunctional organelles (*24*). Proper function of autophagy is essential for preserving cellular proteostasis (*24, 25*). The process can be activated by several stress signals, like amino acid starvation, serum starvation, protein aggregate formation and oxidative stress, and it participates in many processes at the molecular level (recycling of macromolecules and protein aggregate clearance) and the organismal level (bone formation, development and immune function) (*26–35*). Autophagy is regulated by the mammalian target of Rapamycin (mTOR) kinase system (*36*). Under basal conditions, mTOR is active and hinders induction of autophagy; however, under the stresses described above, mTOR is inhibited thereby leading to the activation of autophagy. It should also be noted that autophagy can be induced independent of mTOR (*37*). A key stage in the autophagic process is the formation of the autophagosome, a double-membrane vesicle that engulfs cytoplasmic material and is transported to the lysosome to form the autophagolysosome (AL), where the engulfed cargo is degraded (*38*). The inability of the cell to degrade the engulfed material results in autophagosome/autophagolysosome accumulation, which has been implicated in the etiology of multiple conditions, like AD, diabetes, Huntington’s disease (HD), and Parkinson’s disease (PD) (*18, 39–42*). In addition, general aneuploidy has been associated with disruption of the PN and specific PN molecular events are associated with aneuploidy (*43–51*). These disruptions include limited autophagic degradation, decreased proteosomal activity, diminished protein folding capacity of chaperones and enhanced sensitivity of aneuploidic models to compounds affecting the PN. Since reduced autophagic degradation is present in aneuploidy, this study aims to compare the autophagic process induced by serum starvation (SS) in DS cells and euploid controls.

Using patient-derived cell models from control subjects and individuals with DS, we show that autophagosomal cargo clearance in DS cells after SS is impaired. Our results also demonstrate colocalization of autophagosomal markers with resident proteins of the endosomal trafficking network. Autophagosome fusion with endosomal vesicles suggests the interference of the endosomal pathway in the autophagic process and provides evidence for the observed diminished autophagic flux in DS. Collectively, our results implicate dysfunctional autophagic clearance and subsequently PN disruption as a candidate cellular mechanism for multiple aspects of pathogenesis in DS.

## Materials and methods

### Reagents and antibodies

The following antibodies were purchased from Abcam: p62 (ab56416), LAMP2A (ab18528). The antibody against LC3B (for western blots and immunofluorescence (IF) with p62) was purchased from Novus Biologics (NB100-2220). At this point it has to be mentioned that LC3B antibodies bind to both LC3-I (non-lipidated form) and LC3-II (lipidated form, marker of autophagy induction) proteins. Therefore, in IF experiments LC3-I species cannot be discerned from LC3-II species and the signal obtained regards both protein populations. The secondary goat anti-rabbit IgG antibody was purchased from R&D systems (HAF008). The Alexa Fluor^®^ 488 secondary goat anti-mouse IgG antibody was purchased from Invitrogen (A11001). The following were purchased from Sigma-Aldrich: anti-LC3B (for IF with Rab5, Rab7 and Rab11) (SAB4200361), Chloroquine diphosphate salt (C6628), anti-β-actin (A5441). The following were purchased from Cell Signaling: anti-NBR1 (#9891), anti-Rab5 (*52*), anti-Rab7 (#9367), anti-Rab11 (*53*). Anti-TFEB was purchased from Bethel laboratories (A303-673A). The following were purchased from Jackson ImmunoResearch: Rhodamine Red™ goat anti-mouse IgG secondary antibody for TRITC (115-295-146), Alexa Fluor^®^ 488 goat anti-rabbit IgG secondary antibody for FITC (111-545-144), HRP goat anti-mouse secondary antibody (115-036-003). The Lysotraker Red DND-99 reagent was purchased from Invitrogen (L752). Cathepsin B activity fluorometric assay kit was purchased by Biovision (#K-140). The TFEB Transcription Factor Activity Assay was purchased from Raybiotech (#TFEH-TFEB).

### Cell culture

Eight fibroblast cell lines were obtained from the Coriell Institute for Medical Research (Table 1). Fibroblasts (passage 6 to passage 11) were cultured in Minimum Essential Medium Eagle including L-glutamine and Earle’s salts (Corning, 10-010-CV), with 10% or 15% fetal bovine serum (FBS) (Gibco-A31160601) and 1% non-essential amino acids (Gibco-11140050). For the serum starvation (SS) experiments, the cells were treated with Minimum Essential Medium Eagle including L-glutamine and Earle’s salts (Corning, 10-010-CV) for eight hours, without addition of FBS. Two induced pluripotent cell lines were generated by Dr. David Russel’s laboratory (University of Washington) and were provided by Dr. Christopher Link (University of Colorado, Boulder) in communication with Gretchen Stein. Both iPSC lines originated from the AG06872 fibroblast cell line through reprogramming (*54*). These cells lines are isogenic since the trisomic iPSC cell line spontaneously lost the extra 21^st^ chromosome and became euploid during reprogramming and clone selection. iPSCs were cultured on matrigel-coated plates (Corning, 354277) in mTESR™1 (Stem Cell Technologies-#85851) including 5x Supplement (Stem Cell Technologies, 85852). Neural progenitor cells (NPC) were derived by transformation of iPSC cell lines to embryoid bodies, followed by neural induction. Briefly, iPSCs were plated on 60mm ultra-low attachment dishes (Corning, CLS2361) in STEMdiff™ neural induction medium (Stem Cell Technologies, 05839). After formation of embryoid bodies, neuronal rosettes were selected and replated in matrigel-coated 6-well plates in neural induction medium. After 5 days, neural induction medium was replaced with STEMdiff™ neural progenitor basal medium (Stem Cell Technologies, 05834) and neuronal rosettes were left to expand to NPCs.

**Table 1.** Fibroblast and iPSC lines used.

| Study ID | Coriell ID | Disease status | Sex | Age (year) |
| --- | --- | --- | --- | --- |
| **Fibroblasts** |  |  |  |  |
| CTL1 | AG04392 | CTL | F | Fetal |
| DS1 | AG06872 | DS | F | 1 |
| CTL2 | AG08498 | CTL | M | 1 |
| DS2 | AG06922 | DS | M | 2 |
| CTL3 | AG07095B | CTL | M | 2 |
| DS3 | AG05397 | DS | M | 1 |
| CTL4 | AG04433 | CTL | M | Fetal |
| DS4 | AG07438 | DS | M | 0.75 |
| Tri8 | GM00496 | CTL | M | 2 |
| Tri18 | GM01359 | DS | M | 0.08 |
| **iPSC** |  |  |  |  |
| C3 | AG06872* | CTL | F | 1 |
| C2 | AG06872 | DS | F | 1 |

\* During reprogramming, this clone lost the third copy of chromosome 21; Therefore it is isogenic to the C2 clone.

### Western blotting (WB)

For Western blotting, 20–40 μg of each cell homogenate was separated via SDS-PAGE utilizing a 15% polyacrylamide gel. Proteins were transferred to a nitrocellulose membrane using a Trans-Blot Turbo transfer apparatus (Bio-Rad). Membranes were blocked with 5% nonfat dried milk in TBS-0.1% Tween (TBS-T) for 20 minutes. Primary antibodies were diluted in TBS-T containing 10% Super Block T20 (Thermo Scientific, 37536) at appropriate dilutions (1:500-1:1000) and allowed to bind to membranes overnight at 4°C. Blots were washed (3x) for 10 min in TBS-T, the blot was then incubated with a horseradish peroxidase conjugated secondary antibody at 1:5000 or an Alexa Fluor 488^®^ secondary antibody diluted in TBS-T containing 10% Super Block T20. Clarity Western ECL Substrate (Bio-Rad, 1705060) was used to detect the HRP of the secondary antibody. ChemiDoc MP imaging system and Image Lab software (Bio-Rad) was used to image and quantify blots. These experiments were conducted independently at least twice by utilizing 2-3 technical replicates in each experiment, and the images presented are representative samples.

### Immunofluorescence (IF)

Cells were plated in a 12-well plate containing glass coverslips (1 coverslip per well) at cell density of 40,000 per well and left to adhere and grow overnight. After treatment, cells were fixed using 3.7% (v/v) paraformaldehyde in PBS and permeabilized using 0.1% (v/v) Triton-X 100 in PBS for 12 minutes. Coverslips were blocked at room temperature using a 1:1 mixture of TBS-T and culture medium (EMEM, 15% FBS, 1% NEAA) for 30 minutes. Then incubated with a primary antibody overnight at 4°C and then washed (3x) using TBS-T. Incubation with TRITC-labeled secondary antibody and/or FITC-labeled secondary antibody and DAPI (1μg/ml) followed for 30 minutes at room temperature. Coverslips were washed (3x) in TBS-T and then mounted on slides using VECTASHIELD anti-fade mounting medium (Vector laboratories, H-1000) and SuperMount (BioGenex, NC9742697) and allowed to dry. Cells were imaged using a Nikon TE2000 microscope with a Nikon C1 confocal imaging system. Each coverslip had five to ten different fields imaged and each experiment was conducted in at least two independent trials for a total of 10-20 images per staining combination, treatment and genotype. Analyses of the confocal images were performed as described previously by Orlicky et al (*55*). Briefly, images were converted to TIFF (signal gain was similar for all groups and treatment combinations) and the fluorescence signal of the proteins of interest was quantified in these images and normalized against DAPI by using the 5I Slidebook software program (Intelligent Imaging Innovations, 3I).

### Transmission electron microscopy (TEM)

Cells were plated in a 6-well plate containing glass coverslips (1 coverslip per well) at cell density of 40,000 cells per well and left to grow and adhere overnight. After treatment, the following TEM protocol was followed: Cells were fixed with 2.5% glutaraldehyde for 3 hours, followed by incubation with 1% osmium tetroxide in sodium cacodylate trihydrate buffer (0.05M) phosphate buffer for 1 hour and 1% aqueous uranyl acetate for 30 minutes. Cells were then dehydrated with ethanol and/or propylene oxide at different concentrations and time cycles for a total of 5 hours. For the infiltration, Embed-812 resin mix and/or propylene oxide was used for over 3 days at different concentrations and time cycles. Resin polymerization followed the infiltration step, by 2, 4, 6-Tris (dimethylaminomethyl) phenol (DMP30) addition at the third day for a total of 6 hours. BEEM® capsules for microtomy were inverted on coverslips incubating with Embed-812 and DMP30 and placed in an oven (60°C) for 3 days. Blocks were separated from coverslips using liquid nitrogen to cool, and then heating on a hot plate. Sixty nanometer thin sections were cut on a Leica UC6 ultramicrotome. Images were captured on an FEI T12 Spirit BT (Tecnai) at 100kV. Each sample was imaged ten times (ten different fields) per treatment and genotype. Quantitation of autophagolysosomes (AL) was based on two criteria: a) presence of a double membrane that is a characteristic of autophagosomes and b) presence of lysosomes. Images were randomized, de-identified and at least two independent assessors were blinded to genotype and treatment during AL counting.

### Lysotracker fluorescence intensity measurement

Cells were plated in a 96-well plate at cell density of 10,000/well and left to adhere and grow overnight. Lysotracker Red was added 45 minutes before the end of the SS treatment (8 hours) at a final concentration of 75nM. Red fluorescence (Ex/Em: 577nm/590nm) was measured by using a fluorescent plate reader (Molecular Devices). Lysotracker fluorescence intensity was normalized against protein abundance and is reported as a % of Lysotracker fluorescence intensity of the CTL sample after SS.

### TFEB DNA-binding activity ELISA assay

Nuclear fractions were obtained from cells at basal levels or after SS using the NE-PER nuclear isolation kit (Thermo Scientific-78833), according to the manufacturer’s protocol. TFEB DNA-binding activity ELISA assay was performed according to the manufacturer’s protocol (Raybiotech, #TFEH-TFEB. A SpectraMax 190 microplate reader (Molecular devices) was used to read the absorbance at 450nm. TFEB DNA-binding activity was normalized against protein abundance and is reported as % of TFEB DNA-binding activity of the CTL sample after SS.

### Cathepsin B activity assay

Cathepsin B activity assay was performed according to the manufacturer’s protocol. Briefly, cells were plated in 10 cm plates at cell density of 300,000/plate and left to adhere and recover until the plate achieved 80-90% confluence. After treatment, cells were lysed and protein concentration of the samples was measured by BCA assay. Then, 50ug of protein was added to a black, flat-bottom 96 plate (50ug per well). Reaction buffer and substrate were added in each well and the sample mix was incubated for one hour at room temperature. Negative controls containing a cathepsin-B inhibitor were also used. Fluorescence (Ex/Em: 400nm/505nm) was measured by using a fluorescent plate reader (Molecular Devices). Cathepsin B activity was normalized against protein abundance and is reported as % of Cathepsin B activity of the CTL sample after SS.

### Statistics

Data were analyzed and graphs were plotted using GraphPad Prism 6 software. Data is represented as the mean ± standard error of the mean (SEM). Each experiment was repeated independently at least twice. Statistical significance was determined using unpaired t-test and a P-value of <0.05 was deemed to be significant (* P<0.05; ** P<0.01; *** P<0.001; **** P<0.0001).

## Results

### DS fibroblasts exhibit impaired autophagic degradation after SS

To investigate possible dysfunction of the autophagic process in patient-derived DS cell models, we chose to utilize SS to induce autophagy, since this treatment results in mTOR inhibition and subsequent induction of autophagy (*32, 37*). Our rationale for the use of SS instead of chemical induction (e.g Torin1, Rapamycin) was that it is the least physiologically intrusive method and least molecularly promiscuous strategy to induce autophagy. During autophagy LC3-II, a lipidated form of LC3-I, serves as a marker of successful autophagosome formation since it is a major component of the autophagosome membrane (*56*). Similarly, the engulfed cargo includes proteins of the autophagy receptor family, with p62 and NBR1 serving as prominent members (*57*). These receptor proteins bind ubiquitinated macromolecules and form the core of the autophagosomal cargo destined for degradation (*58*). Impaired clearance of autophagic cargo results in autophagosome/autophagolysosome accumulation.

To examine any basal or SS-mediated differences in the relative amount of AL and to compare autophagic degradation efficiency in DS cells, we employed TEM analyses in a DS and euploid CTL fibroblast cell line. These experiments revealed an increased number of AL in a DS fibroblast cell line at untreated conditions compared to euploid CTL cell (Figure 1A-B). However, TEM analyses typically utilize a limited set of fields and different research groups base their analyses on different criteria to label AL or other cellular entities. Therefore, we next chose to evaluate autophagy induction and completion via WB analyses of LC3-II and p62 and to extend these analyses by inclusion of a larger number of fibroblast pairs. Western blotting of four CTL and DS fibroblast lines showed no significant difference in basal LC3-II or p62 abundance (Figure 1C-D). These data indicate that DS fibroblasts do not differ compared to disomic CTL in markers of basal autophagy. When the same cell lines were treated with SS, we observed a marked increase in the number of AL in DS cells compared to euploid CTL cells via TEM analysis (Figure 1A-B). Increased presence of AL is a marker of impaired autophagic flux in DS fibroblasts and is indicative of the impaired ability to successfully degrade autophagosomes in trisomic cells (*59*).

Consistent with the TEM results, IF for p62 revealed significantly increased levels of this autophagy receptor in a DS fibroblast cell line after SS compared to the euploid CTL group (Figure 1E-F). Impeded degradation of p62 after autophagic induction in DS cells is a marker of dysfunctional autophagy. The expression levels and subcellular location of p62 are not exclusively regulated by autophagy and this protein can be impacted by other regulatory pathways, like the Nrf2-Keap-1 system (*60–62*)). In order to confirm p62 accumulation is due to impaired general autophagic clearance and not just an effect of p62 metabolism, we evaluated an additional autophagy receptor, NBR1. Interestingly, IF for NBR1 demonstrated that this protein follows a similar pattern to p62 with a significant increase in abundance in the DS fibroblast line after SS compared to diploid CTL (Supplemental Figure 1). This result indicates that impaired clearance is not specific to p62 since other autophagy receptors exhibit the same pattern. Taken together, increased abundance of AL, p62 and NBR1 after SS in DS cells indicates significantly impeded autophagic degradation in this condition.

**Figure 1.**
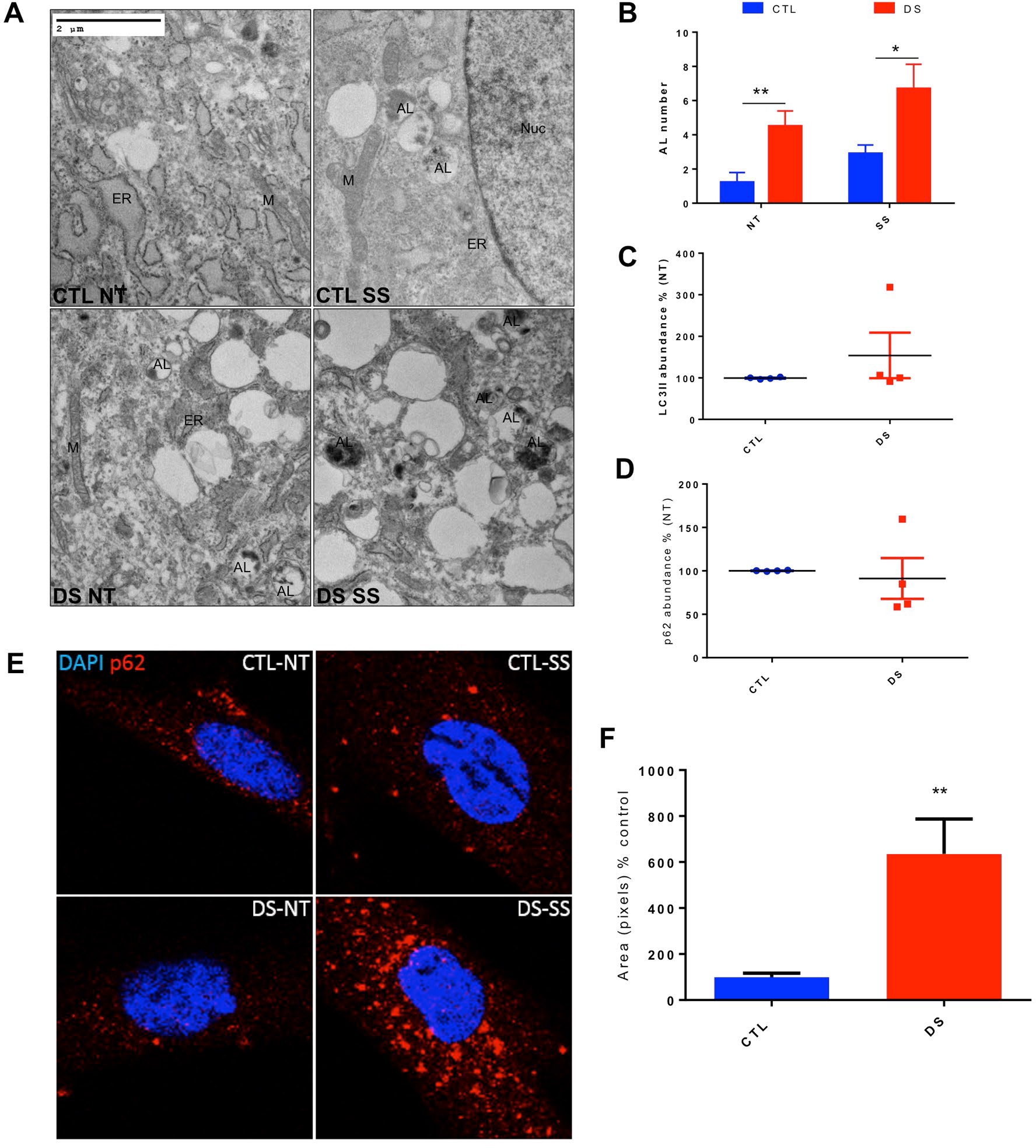
DS fibroblasts exhibit increased number of autophagolysosomes and autophagosome cargo abundance after SS. (A) TEM figures of a CTL and DS fibroblast cell line at basal levels or after SS. (B) Quantitation of autophagolysosome (AL) number of TEM figures at basal levels. (C) Quantification of abundance levels of LC3-II of four cell line pairs of CTL and DS fibroblasts at basal levels (% based on abundance of CTL NT, representative blot in Figure 2A). (D) Quantification of % abundance levels of p62 of four cell line pairs of CTL and DS fibroblasts at basal levels (% based on abundance of CTL NT, representative blot in Figure 2A). (E) Immunofluorescence for p62 (Red) and DAPI (blue) at basal levels or after SS. (F) Quantification of p62 fluorescence intensity of the immunofluorescence experiment (% based on abundance of CTL SS). ER, endoplasmic reticulum; M, mitochondria; NT, Not treated-basal levels; Nuc, Nucleus; SS, serum starvation (8h).

### Autophagosome accumulation is induced by SS in DS fibroblasts

To further investigate the results described above, we next used Western blot analyses to measure SS-mediated alterations in LC3-II and p62 protein levels in four pairs of CTL and DS fibroblasts. Following SS, the scale of induction of LC3-II was essentially identical between CTL and DS cell lines (Figure 2A&B), indicating successful mTOR inhibition and autophagic activation in both groups. Consistent with the results described above (Figure 1), the scale of SS-mediated increase of p62 abundance was significantly elevated in DS cells relative to euploid controls (Figure 2A&C). Successful induction of autophagy (increase in LC3-II levels) and dysfunction of the degradation step (p62 accumulation) were also observed in a DS iPSC cell line and a DS NPC cell line (Supplemental Figure 2). These results are consistent with our hypothesis that limited autophagic degradation (p62 accumulation after SS) and subsequent PN dysfunction are features of aneuploidic cells and can occur in DS cells at several stages of differentiation. Furthermore, Western blot analyses of p62 abundance at several different time points following SS revealed that DS fibroblasts accumulate p62 by 8h and that this increase in protein level persists even after 24h of SS (Supplemental Figure 3). These results further demonstrate that successful degradation of autophagosomal cargo in DS cells is impaired.

**Figure 2.**
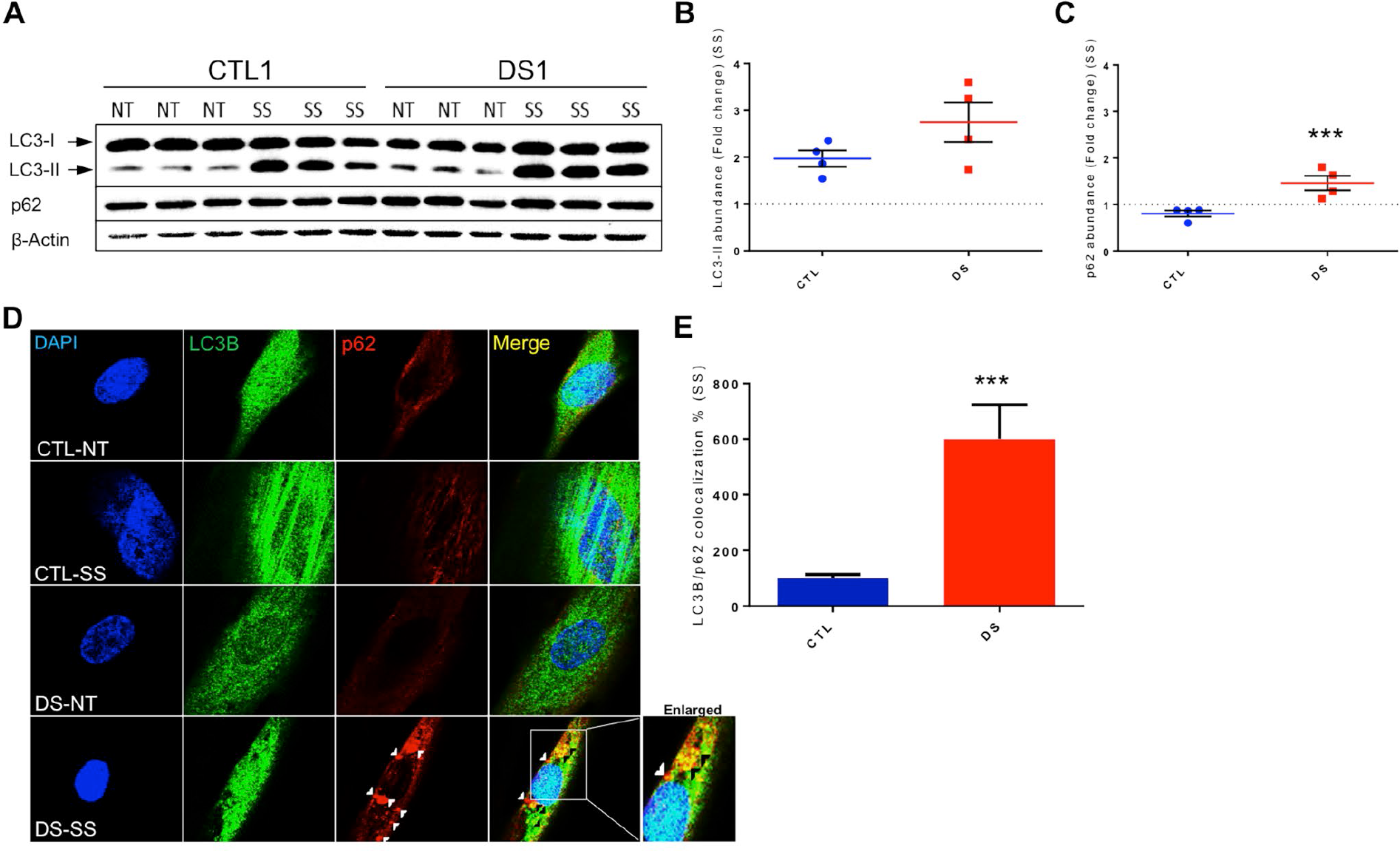
DS fibroblasts display autophagosome accumulation and increased p62 abundance after SS. (A) Representative blot of LC3-I, LC3-II, p62 and β-actin of CTL and DS fibroblasts (three technical replicates per treatment and genotype are presented). (B) Quantification of fold change in abundance of LC3II of 4 cell line pairs of CTL and DS fibroblasts after SS. (C) Quantification of fold change in abundance of p62 of four cell line pairs of CTL and DS fibroblasts after SS. (D) IF for LC3B (Green), p62 (Red) and DAPI (Blue) in a CTL and DS fibroblast cell line at basal levels or after SS. (E) Quantification of LC3B/p62 co-localization (yellow) fluorescence intensity (area-pixels) after SS (% based on abundance of CTL SS). White arrowheads= p62 only puncta Black arrowhead - p62/LC3B co-localized puncta. NT, Not treated-basal levels; SS, serum starvation (8h).

To expand our investigation of the autophagic process and to determine if p62 can be successfully engulfed into autophagosomes, IF for p62 and LC3B was performed following SS. These experiments showed partial co-localization of these two proteins in DS fibroblasts after SS (Figure 2D-E), which demonstrates successful cargo engulfment by the autophagosome. The co-localization of p62 and LC3B also indicates the presence of undegraded autophagosomes in DS fibroblasts after SS. This result further supports our hypothesis that DS cells exhibit a dysfunctional PN and inefficient autophagic clearance.

### Trisomy 8, but not trisomy 18, fibroblasts also display impairment of autophagic clearance after SS

Limited autophagic degradation is observed in models of aneuploidy (*47*). In our research, it is presently unknown if this particular phenotype is a function of specific genes on chromosome 21 or simply a generalized consequence of aneuploidy. To evaluate possible dysfunction of autophagy, LC3-II and p62 protein abundance were examined via WB in a trisomy 8 (Tri8), trisomy 18 (Tri18) and CTL fibroblast cell line after SS. Both trisomic cell lines exhibited successful induction of autophagy since LC3-II abundance was increased after SS compared to basal levels (6A&C). The scale of induction observed was similar in both trisomic lines and was consistent with our previous results in DS fibroblasts (Figure 2). Similar to what we have observed in DS fibroblasts, Tri8 fibroblasts also displayed significantly higher p62 accumulation after SS compared to CTL cell models (Figure 3A&B). However, results from Tri18 fibroblasts showed no difference in p62 abundance after SS compared to CTL cell models, indicating successful autophagic degradation in this aneuploidic condition. Collectively, these data indicate that autophagic dysfunction can occur in different forms of trisomy but is not necessarily a generalized phenomenon of aneuploidy. Therefore, each chromosomal duplication should be investigated independently in order to determine alternative hallmarks of general PN dysfunction in each type of trisomy.

**Figure 3.**
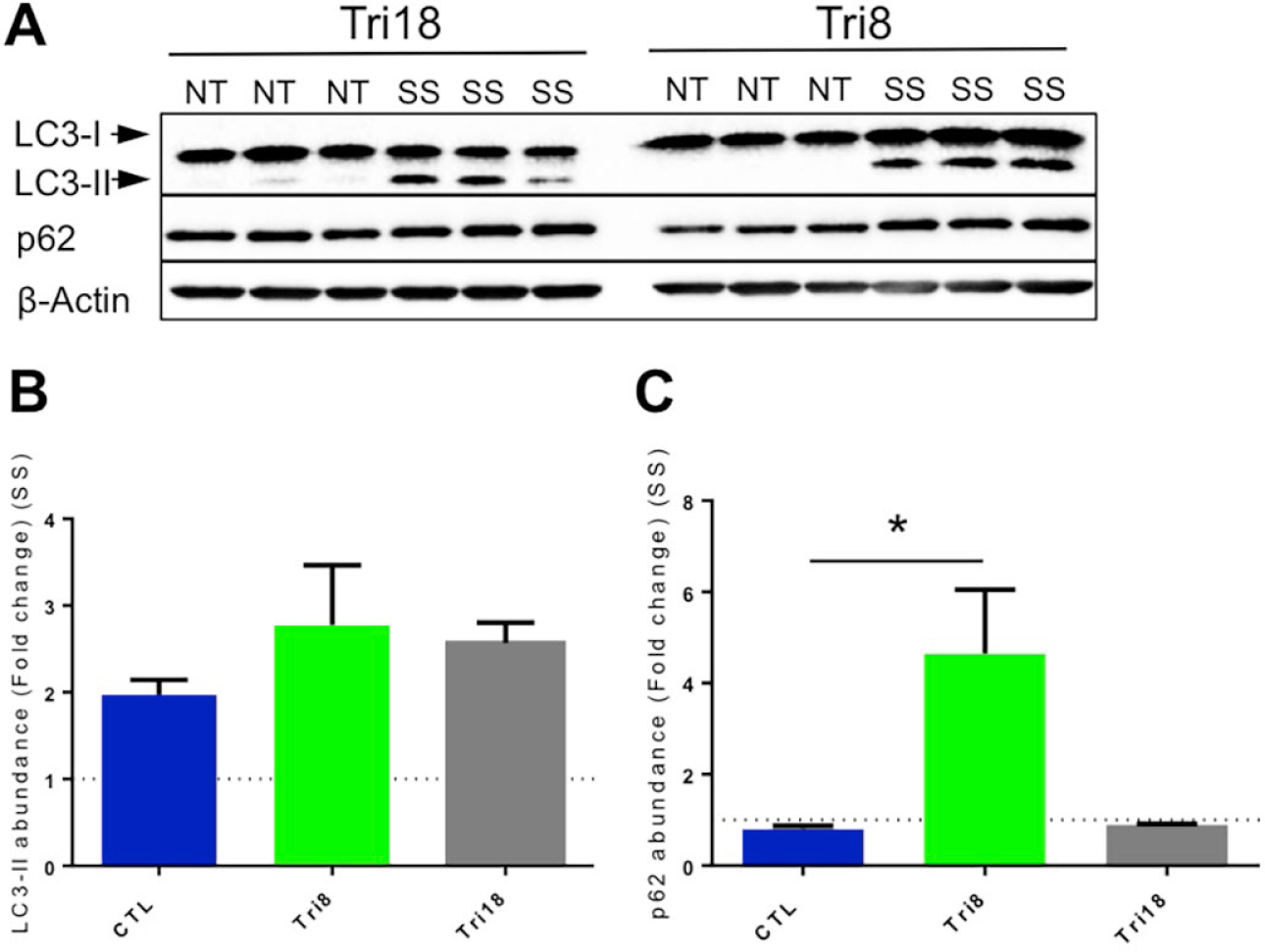
p62 accumulation after SS is also observed in Tri8 fibroblasts, but is absent in Tri18 fibroblasts. (A) Representative blot of LC3I, LC3II, p62 and β-actin of Tri8 and Tri18 fibroblast cell lines at basal levels or after SS (three technical replicates per treatment and genotype are presented). (B) Quantification of fold change in abundance of LC3II of CTL (reported also in Figure 2A), Tri8 and Tri18 fibroblasts. (C) Quantification of fold change in abundance of p62 of CTL (reported also in Figure 2A), Tri8 and Tri18 fibroblasts. NT=Not treated-basal levels SS=serum starvation (8h).

### Lysosomal dysfunction is not responsible for p62 accumulation in DS fibroblasts after SS

Motivated by the observation that p62 levels remain elevated after SS in DS cells and that the lysosome is critically important for autophagic cargo degradation (*63, 64*), we next evaluated lysosomal endpoints to examine AL accumulation and possible lysosomal dysfunction in DS fibroblasts after SS. For these assessments, CTL and DS fibroblasts were co-treated with the lysosomal inhibitor chloroquine (CQ) and SS. CQ is a well-characterized inhibitor of lysosomal proteolytic activity because it alters lysosomal pH and renders lysosomal resident proteases inactive (reviewed here (*65*)). Therefore, if lysosomal dysfunction is responsible for the observed impairment of autophagic degradation in DS cells, SS co-treatment with CQ or SS alone should display similar levels of p62 and LC3-II in these DS cells. Although not statistically significant, we observed that CQ and SS co-treatment resulted in apparent increased abundance of p62 and LC3-II compared to SS treatment alone (Figure 4A-C). The observation that p62 and LC3-II protein levels were decreased in the absence of CQ indicates that the lysosome is unlikely to be the source of the observed impairment of autophagic degradation.

We further assessed autophagosome-lysosome fusion via IF by investigating co-localization of p62 and the lysosome-associated membrane protein 2A (LAMP2A), a lysosomal transmembrane protein (*66*). LAMP2A is ubiquitously expressed in the lysosomes, it is a well characterized marker of lysosomal localization and is thought to promote membrane integrity for lysosomal stability (*67*). Our experiments showed that SS lead to partial co-localization of p62 with LAMP2A in DS cells (Figure 4D-E), further supporting the AL accumulation observed in our TEM experiment. Presence of “p62-only” puncta and only partial p62 and LAMP2 co-localization in DS cells is also consistent with the possibility that disrupted autophagosome-lysosome fusion or sequestration of autophagosomes in other locations within the cell may be responsible for the impairment of autophagic degradation.

To expand our examination of the effect of SS on lysosomes in both CTL and DS, Lysotracker dye was utilized. Lysotraker is a lysomotropic dye that emits fluorescence when entering acidic organelles, like lysosomes (*68*). Lysotracker fluorescence intensity was measured in CTL and DS fibroblast cell lines after SS, to investigate possible lysosomal pH disturbance. No significant difference in intensity was observed in the CTL or DS fibroblasts, indicating normal pH levels between the two groups (Figure 4F). Additionally, the lysosome contains proteolytic enzymes called cathepsins (*63, 64*). These enzymes are functional only in the lysosomal space due to optimal acidic pH conditions. In our experiment, we evaluated cathepsin B function as an index of lysosomal degradation capacity since inhibition of activity may explain dysfunctional clearance in DS fibroblasts. Remarkably, cathepsin B activity did not differ significantly between CTL and DS cells after SS, indicating that the lysosome is not overtly dysfunctional in DS (Figure 4G).

**Figure 4.**
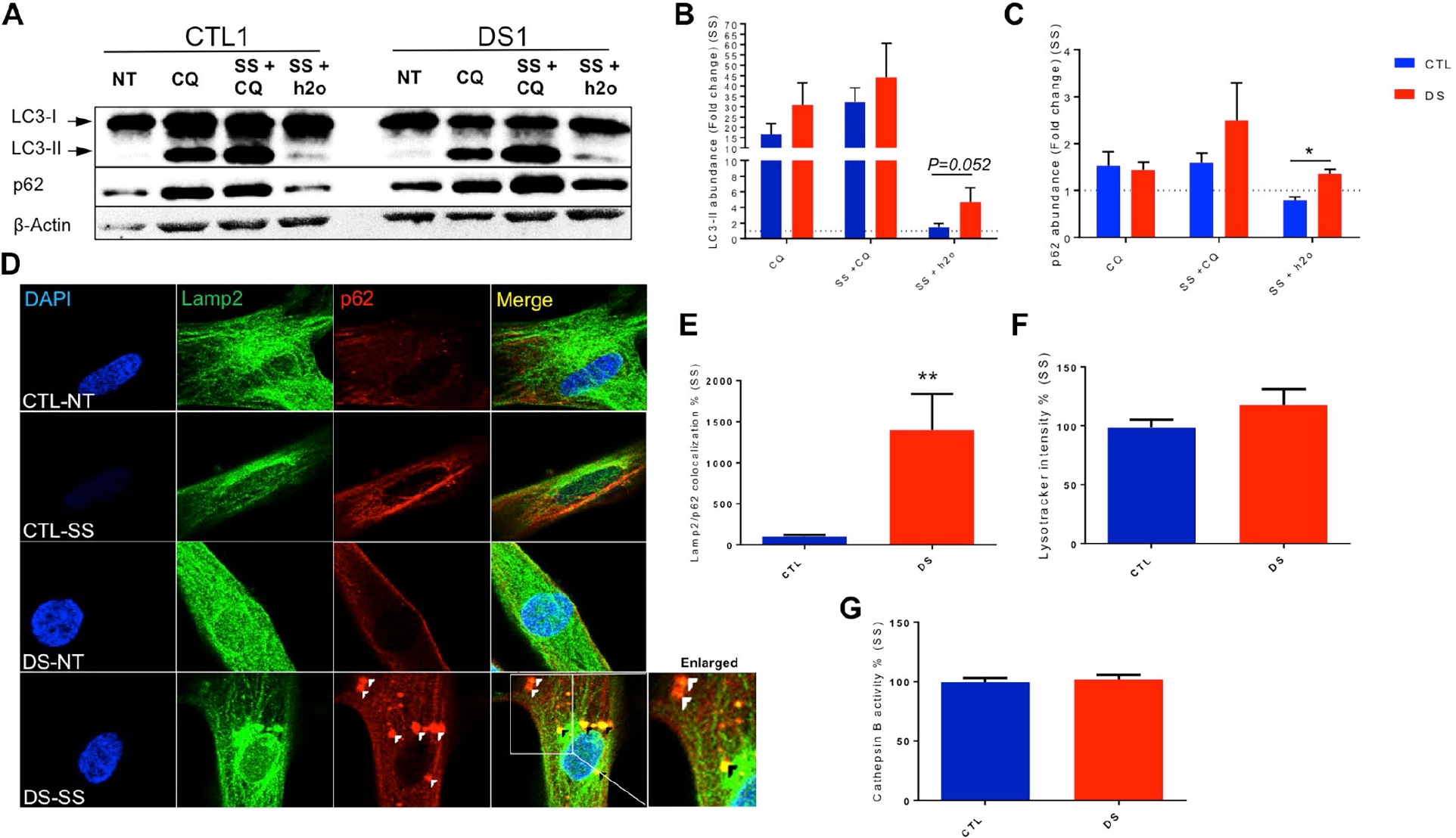
Lysosomal dysfunction is not responsible for the observed p62 accumulation. (A) Representative blot of LC3-I, LC3-II, p62 and β-actin of a CTL and DS fibroblast cell line at basal levels, after CQ treatment, after SS and CQ co-treatment and after SS and h2o co-treatment used as control. (B) Quantification of fold change in abundance of LC3-II of a CTL and DS fibroblast cell line after CQ, SS+CQ and SS+h2o. (C) Quantification of fold change in abundance of p62 of a CTL and DS fibroblast cell line after CQ, SS+CQ and SS+h2o. (D) IF for LAMP2 (Green), p62 (Red) and DAPI (Blue) in a CTL and DS fibroblast cell line at basal levels or after SS. (E) Quantification of LAMP2/p62 co-localization (yellow) fluorescence intensity (area-pixels) after SS (% based on abundance of CTL SS). (F) Lysotracker-Red fluorescence intensity in a CTL and DS fibroblast cell line after SS (% based on fluorescence intensity of CTL SS). (G) Proteolytic activity of lysosomal cathepsin B (% based on activity of CTL SS), White arrowhead - p62 only puncta; Black arrowhead - p62/LAMP2 co-localized puncta.

TFEB is a transcription factor responsible for regulating lysosomal biogenesis (*69*). At basal conditions, TFEB is sequestered by mTOR on the lysosomal membrane surface where it is kept in an inactive state. Inhibition of mTOR by SS results in TFEB translocation to the nucleus where it promotes the expression of a battery of genes involved in lysosomal biogenesis. We further examined lysosomal dysfunction via assessment of activation of TFEB after SS by using a combination of IF and TFEB DNA-binding activity ELISA. Similar results regarding nuclear localization and DNA binding of TFEB were observed in both CTL and DS fibroblasts after SS (Supplemental Figure 4), again indicating that the defects in autophagic degradation in DS models after SS may not be explained by simple disruption of lysosomal physiology and signaling.

### Autophagosome markers colocalize with early endosomes (Rab5) in DS cells after SS

Previous research has indicated that the autophagic process is subject to significant regulatory interaction with the endosomal trafficking system, which is heavily regulated by members of the Rab protein family(*70*). Also, DS models have been reported to exhibit enlarged early endosomes and increased endocytic uptake (*71, 72*). Therefore, we next evaluated possible cross-talk between autophagy and the endosomal trafficking pathway. The early endosome serves as sorting station for endocytosed cargo and is the first organelle of the endocytic machinery to receive incoming material. Rab5, a prominent member of the Ras GTPase family, resides in early endosomal vesicles (*70, 73*) and is involved in endocytosis and endosomal sorting (*74*) In order to examine possible interaction between autophagic vesicles and early endosome function in DS, we investigated the effect of SS on Rab5 protein abundance in DS fibroblasts relative to euploid controls. Basal abundance of Rab5 was observed to be similar between CTL and DS fibroblasts (Figure 5A&B). Additionally, Rab5 abundance after SS did not differ significantly between the CTL and DS fibroblast cell lines (Figure 5A&C). Interestingly, combined IF for both Rab5 and p62 or Rab5 and LC3B revealed increased Rab5/LC3B and Rab5/p62 co-localization in DS fibroblasts compared to CTL after SS (Figure 5D-G). These results indicate possible fusion of autophagosomes with early endosomes, which might impede both endocytic trafficking and autophagic degradation.

**Figure 5.**
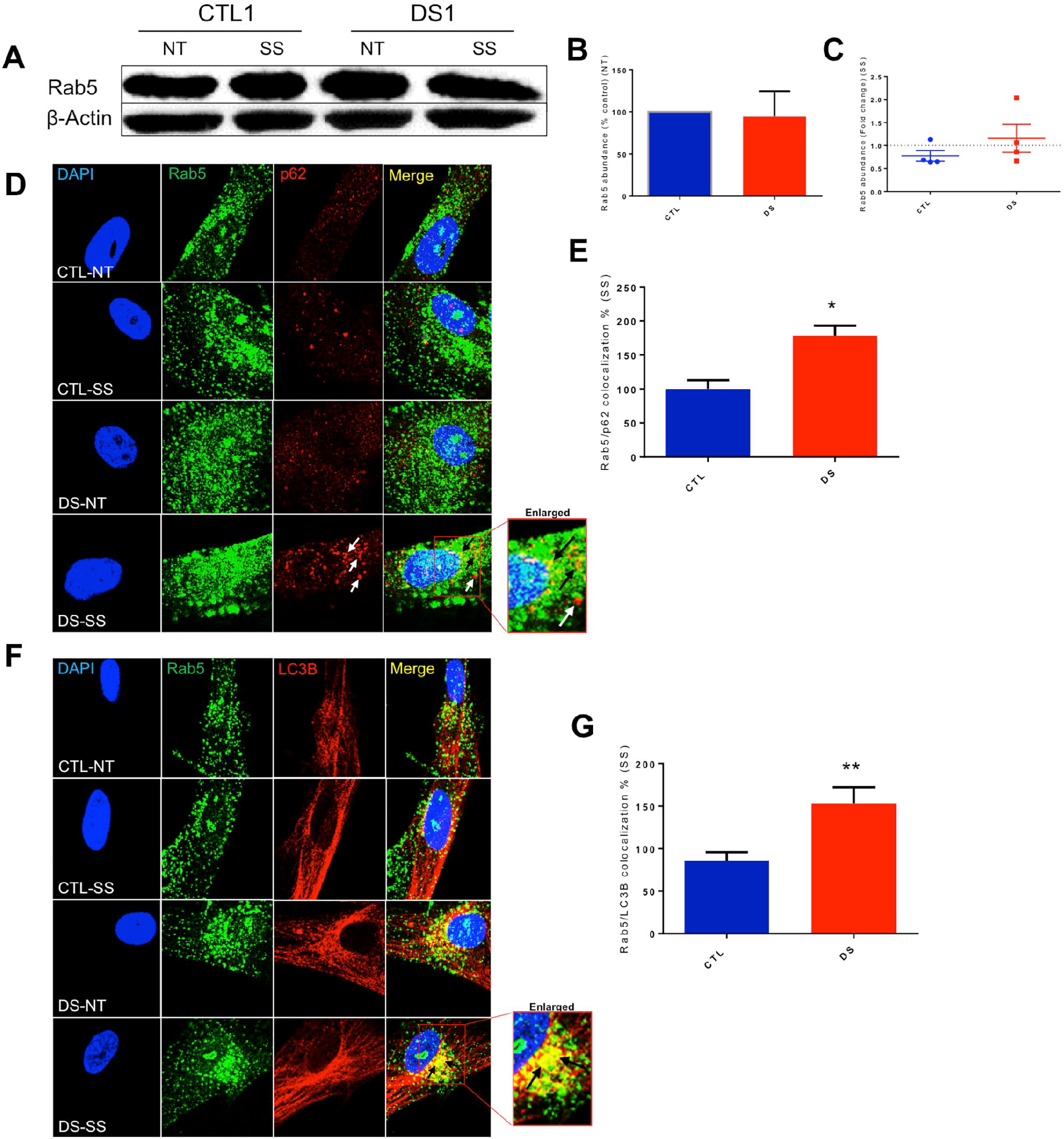
DS fibroblasts exhibit increased p62 and LC3B co-localization with the early endosome marker Rab5 after SS treatment. (A) Representative blot of Rab5 and β-actin of CTL and DS fibroblast cell lines at basal levels or after SS. (B) Quantification of % abundance levels of Rab5 of four cell line pairs of CTL and DS fibroblasts at basal levels (% based on abundance of CTL NT). (C) Quantification of fold change in abundance of Rab5 of a CTL and DS fibroblast cell lines after SS. (D) IF for Rab5 (Green), p62 (Red) and DAPI (Blue) in a CTL and DS fibroblast cell line at basal levels or after SS. (E) Quantification of Rab5 (green) and Rab5/p62 co-localization (yellow) fluorescence intensity (area-pixels) after SS (% based on abundance of CTL SS). (F) IF for Rab5 (Green), LC3B (Red) and DAPI (Blue) in a CTL and DS fibroblast cell line at basal levels or after SS. (G) Quantification of Rab5 (green) and Rab5/LC3B co-localization (yellow) fluorescence intensity (area-pixels) after SS (% based on abundance of CTL SS). White arrow - p62 only puncta; Black arrow - Rab5/p62 or Rab5/LC3B colocalized puncta; NT - Not treated-basal levels; SS - serum starvation (8h).

### Late endosomes (Rab7) colocalize with autophagosome markers in DS cells after SS

Driven by our results regarding Rab5, we next evaluated the effect of SS on late endosomes in DS fibroblasts. A key event during endocytic trafficking is the maturation of early endosomes to late endosomes by replacement of Rab5 with Rab7 (*75, 76*). Similar to the Rab5 results, co-localization of late endosome markers with autophagosome proteins is possible, since direct late endosome fusion with autophagosomes has been reported (*77, 78*). Rab7 is a critical regulator of endocytic trafficking, is responsible for cargo transportation to the lysosome for degradation, and regulates fusion of late endosomes with the lysosome (*79*). Basal abundance of Rab7 was also observed to be similar between CTL and DS fibroblasts (Figure 6A&B). No significant difference between the CTL and DS fibroblast cell lines was observed regarding Rab7 induction after SS (Figure 6A&C). However, similar to the results regarding Rab5, IF for Rab7 and p62 or Rab7 and LC3B revealed increased Rab7/LC3B and Rab7/p62 co-localization in DS fibroblasts compared to CTL after SS (Figure 6D-G). Co-localization of autophagosome markers with both early endosomes and late endosomes suggests disruption of total endosomal trafficking in DS during SS-mediated autophagy, rather than simple sequestration of autophagosomes in specific endocytic compartments that could prevent their degradation. Also, since late endosomes can fuse directly with the lysosome (*80, 81*), fusion of autophagosomes with Rab7-positive vesicles might result in abnormal levels of cargo destined for degradation in the lysosome and subsequently impair autophagic clearance. This mechanism could explain AL accumulation in DS fibroblasts in the absence of lysosomal defects.

**Figure 6.**
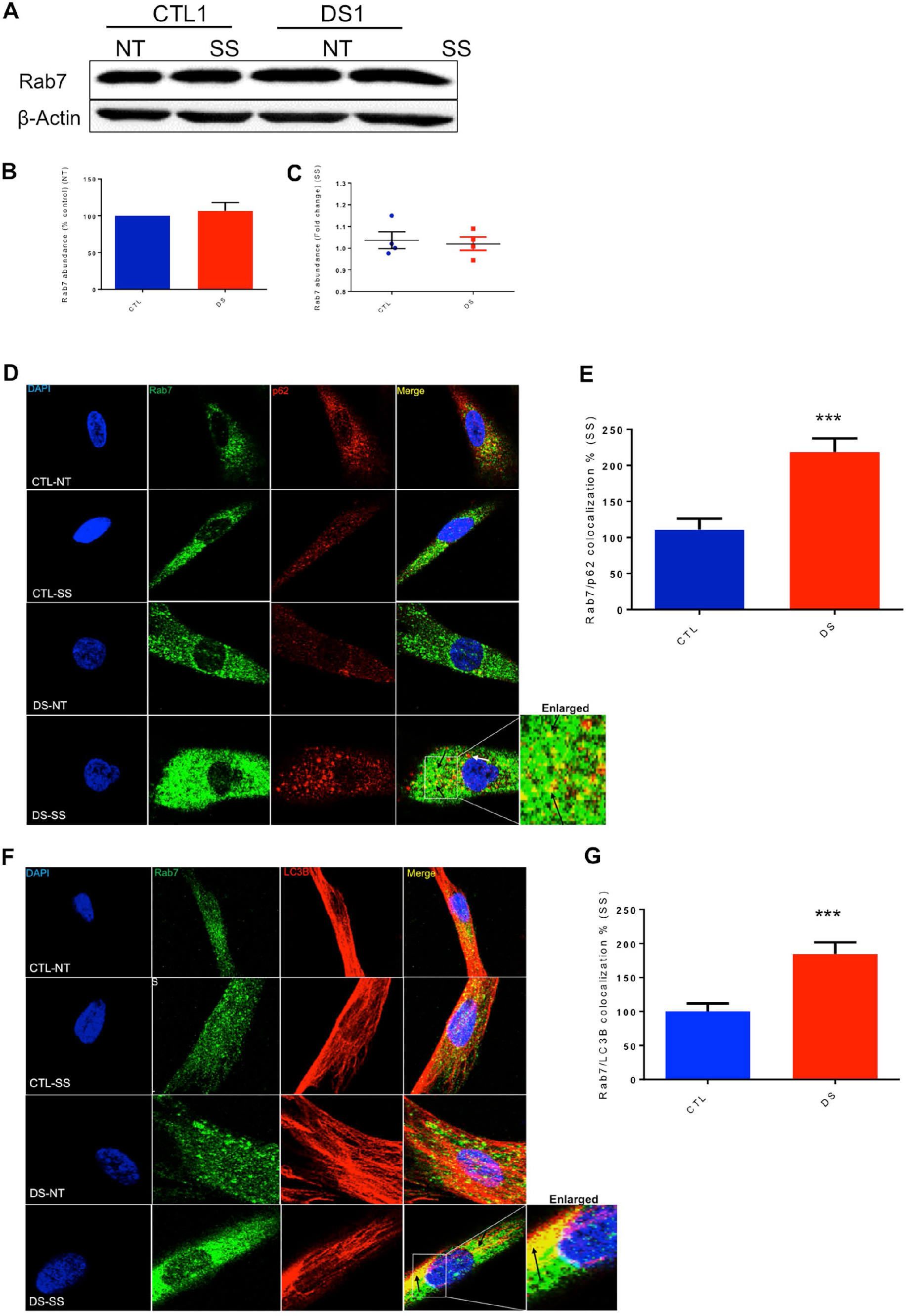
DS fibroblasts exhibit increased p62 and LC3B co-localization with the late endosome marker Rab7 after SS treatment. (A) Representative blot of Rab7 and β-actin of CTL and DS fibroblast cell lines at basal levels or after SS. (B) Quantification of % abundance levels of Rab7 of four cell line pairs of CTL and DS fibroblasts at basal levels (% based on abundance of CTL NT). (C) Quantification of fold change in abundance of Rab7 of a CTL and DS fibroblast cell lines after SS. (D) IF for Rab7 (Green), p62 (Red) and DAPI (Blue) in a CTL and DS fibroblast cell line at basal levels or after SS. (E) Quantification of Rab7 (green) and Rab7/p62 co-localization (yellow) fluorescence intensity (area-pixels) after SS (% based on abundance of CTL SS). (F) IF for Rab7 (Green), LC3B (Red) and DAPI (Blue) in a CTL and DS fibroblast cell line at basal levels or after SS. (G) Quantification of Rab7 (green) and Rab7/LC3B co-localization (yellow) fluorescence intensity (area-pixels) after SS (% based on abundance of CTL SS). White arrow - p62 only puncta; Black arrow - Rab7/p62 or Rab7/LC3B co-localized puncta; NT - Not treated-basal levels; SS - serum starvation (8h).

### Rab11 exhibits significantly increased abundance and colocalization with autophagosome markers in DS fibroblasts

In an effort to expand our research by studying other Rab-family members, we investigated possible involvement of a recycling endosome marker, Rab11, in the autophagic process induced by SS. This protein can function as a signaling nexus between the endosomal pathway and autophagy (*82, 83*). In our experiment, the scale of increase in Rab11 abundance was significantly higher in our four DS fibroblast cell lines after SS compared to diploid CTL (Figure 7A&C). In addition, Rab11 accumulated in DS fibroblasts in a manner similar to p62, which suggests that recycling endosome accumulation takes place in DS fibroblasts after SS. Moreover, increased Rab11/LC3B and Rab11/p62 co-localization was also observed after SS (Figure 7D-G), further supporting the results obtained in the experiments regarding Rab5 and Rab7 (Figure 5, Figure 6). This observed autophagosome fusion with Rab11-positive vesicles indicates possible inhibition of the recycling endocytic pathway that results in Rab11 accumulation. Also, since autophagosomes fuse with many types of endosomal vesicles (early, late and recycling), inhibition/collapse of total endosomal trafficking is suggested.

**Figure 7.**
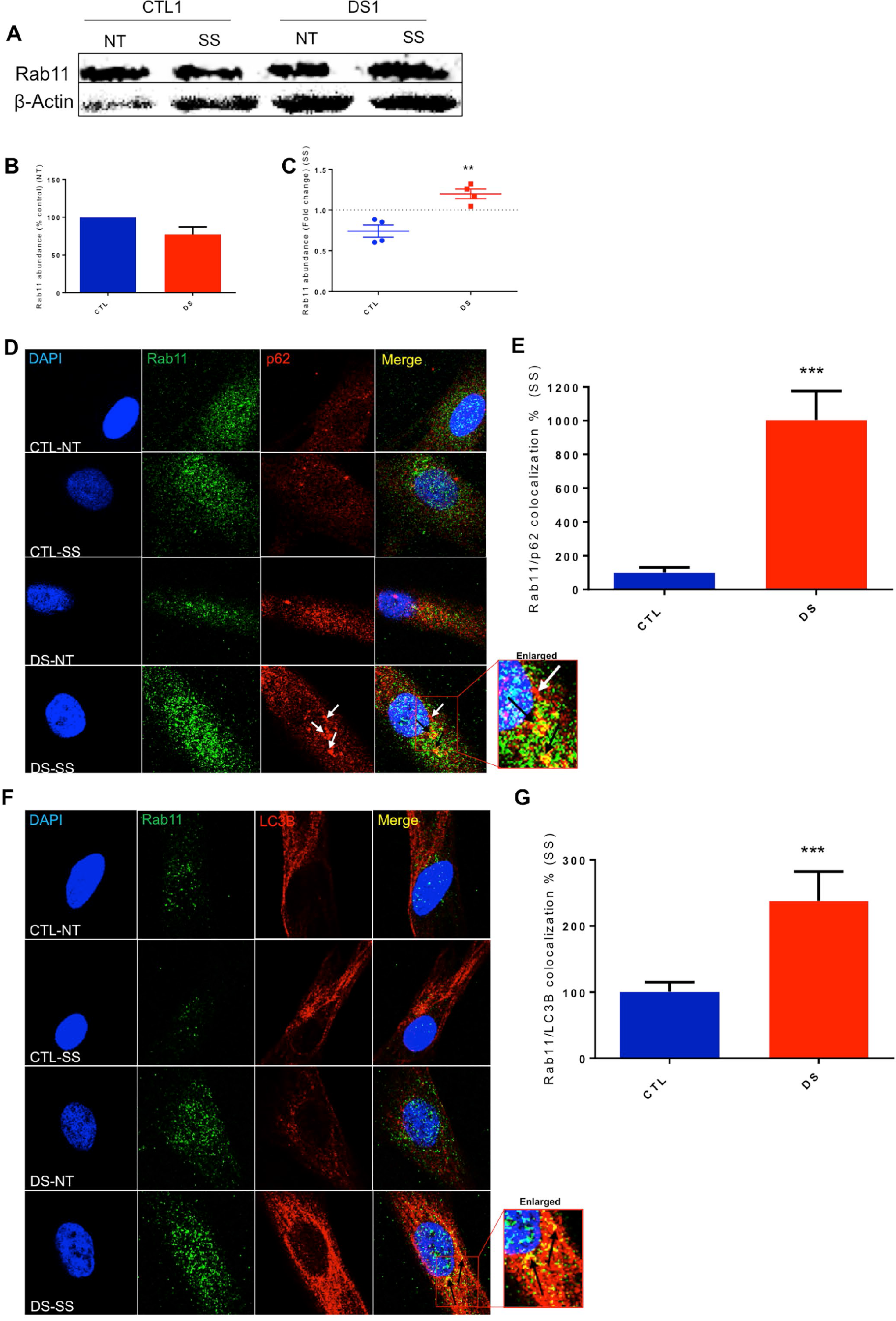
Increased colocalization of Rab11 with p62 and LC3B is observed in DS fibroblasts after SS. (A) Representative blot of Rab11 and β-actin of CTL and DS fibroblast cell lines at basal levels or after SS. (B) Quantification of % abundance levels of Rab11 of four cell line pairs of CTL and DS fibroblasts at basal levels (% based on abundance of CTL NT). (C) Quantification of fold change in abundance of Rab11 of a CTL and DS fibroblast cell lines after SS. (D) IF for Rab11 (Green), p62 (Red) and DAPI (Blue) in a CTL and DS fibroblast cell line at basal levels or after SS. (E) Quantification of Rab11 (green) and Rab11/p62 co-localization (yellow) fluorescence intensity (area-pixels) after SS (% based on abundance of CTL SS). (F) IF for Rab11 (Green), LC3B (Red) and DAPI (Blue) in a CTL and DS fibroblast cell line at basal levels or after SS. (G) Quantification of Rab11 (green) and Rab11/LC3B co-localization (yellow) fluorescence intensity (area-pixels) after SS (% based on abundance of CTL SS). White arrow - p62 only puncta; Black arrow - Rab11/p62 or Rab11/LC3B co-localized puncta; NT - Not treated-basal levels; SS - serum starvation (8h).

## Discussion

The presence of an extra copy of chromosome 21 in the DS genome puts DS individuals at high risk for developing certain comorbidities, such as AD, immune dysfunction, diabetes and leukemia (*1, 4, 6, 7, 84–86*). Many of the DS comorbidities are associated with a dysfunctional PN (*14–21*) and aneuploidy has also been associated with proteotoxic stress and PN dysfunction (*43–51*). Furthermore, PN collapse is observed in aging and DS can be considered a condition of accelerated senescence and aging (*87–92*). These findings inspired our interest in investigating possible PN dysfunction and the presence of a dysregulated protein quality control system in DS. Recently, our laboratory has reported a dysfunctional PN in DS models with increased presence of ER stress, limited chaperone expression after heat stress and increased sensitivity to proteotoxic compounds (*8*). Our findings here build upon this previous research and focus on further characterizing the possible pathogenic role of PN disruption via the study of the autophagic process in DS models.

To date, research regarding the autophagic process in DS has been limited; however, two independent groups have reported mTOR hyperactivation in postmortem human DS brain samples (*9, 93*). Hyperactivity of mTOR in DS human samples might be an indication of general dysregulation of the mTOR pathway that promotes dysfunction of general autophagy in the DS phenotype. However, this data should be treated with some caution since medication and pathologies that might affect mTOR function and the autophagic pathway are unknown in these postmortem samples. Therefore, our approach using primary cells, like patient-derived fibroblasts, provides the advantage of removing confounding factors and constitutes a more stringently controlled system to evaluate dynamics of the autophagic process. Furthermore, the use of a cell culture model and SS-mediated induction of autophagy utilized here was intended to specifically focus on mechanisms that occur downstream of mTOR inhibition.

Our initial TEM results revealed an increased presence of AL at basal levels in DS fibroblasts, which is a similar result to the previously reported presence of accumulated autophagosomal proteins within lysosomes in chromosome missegregation models (*47*). However, Western blot results showed that DS and euploid control cells possess similar basal levels of LC3-II and p62, indicating that DS cells do not exhibit impaired autophagic flux at basal levels. These data indicated a necessity to investigate stimulus-induced autophagy to confirm or deny if this key process is dysfunctional in a model that already displays markers of altered proteostasis. Successful induction of autophagy after SS was observed in all DS-individual derived cell models, since no difference regarding LC3-II protein abundance was observed in DS compared to the CTL group. In addition, SS resulted in diminished autophagy receptor (p62, NBR1) degradation in DS cell models, indicating that impaired autophagic degradation occurs in DS. This impairment in autophagic degradation has also been observed in AD, PD, HD and other neurological disorders (*21, 94–97*). Additionally, the observation of impeded autophagic clearance in several DS cell models at different stages of differentiation (fibroblasts, iPSCs, NPCs) also suggests that this impairment may occur *in utero* and in a relatively wide range of cell types. The significant persistence of dysfunctional protein degradation throughout our experiments provides a rationale for labeling impaired autophagic clearance as a cellular phenotype of DS. Further investigation is needed, however, to define possible molecular signatures of autophagic dysfunction in DS, such as localization and relative kinetics of the various molecular events involved. To this end, cell models are appropriate and informative experimental tools for this task and the possible impairment of autophagic clearance through mTOR-independent pathways in DS is currently the subject of investigation in our laboratory.

A similar model of inhibited autophagosomal clearance has been reported in alternative aneuploidic models, although this event occured in cells at basal levels only and not after SS (*47*). These particular results would indicate that dysfunctional clearance may be a generalized feature of chromosomal missegregation (as proposed in (*47*)) but also highlights potential differences between each type of aneuploidy. Poor p62 clearance observed in Tri8 fibroblasts described here, but not in Tri18 fibroblasts, supports this argument. Furthermore, aneuploidy is also a common feature of cancer (*44, 98*) and DS individuals have increased risk of developing multiple forms of leukemia, especially during childhood (*6, 7, 84, 99*). However, DS individuals are paradoxically resistant to solid tumor development compared to other aneuploidy models (*5, 84, 100*), confirming alterations in pathological outcomes between different aneuploidies. The effects of general aneuploidy on cellular physiology should be discerned from the effects of individual types of aneuploidy (e.g. Trisomy 21, Monosomy X (*101, 102*)). Lastly, increased life expectancy of DS individuals versus individuals with other trisomies (e.g. Tri18) also supports this conclusion (*2*).

Spatial distribution of p62 and NBR1 accumulation within the cells can shed light on the mechanistic source of clearance inhibition. For example, in our studies p62 and NBR1 accumulation appeared throughout the cytoplasm and was not restricted to a specific subcellular compartment or cellular region. Perinuclear accumulation of autophagy receptors would indicate localization of p62 to lysosomes that reside around the nucleus and is a common observation in cases of dysfunctional autophagic clearance (*103, 104*). Co-localization of p62 with LC3B in DS fibroblasts after SS confirms that autophagosome accumulation takes place, while co-localization of p62 with LAMP2 indicates successful fusion of autophagosomes and lysosomes and corroborates AL accumulation observed in the TEM experiment. However, the presence of “p62-only” puncta is suggestive that p62 may be sequestered in other cellular entities, such as endosomal vesicles or other organelles and explains p62 and NBR1 localization throughout the cell. Likewise, the observation that SS co-treatment with CQ (SS+CQ) resulted in an increased abundance of both p62 and LC3-II in DS compared to the SS-only treatment indicates that the lysosome is probably not responsible for the inhibition of autophagic clearance. Adding further weight to this conclusion are the observations of normal lysosomal pH levels, cathepsin B activity and TFEB activation after SS in both CTL and DS cells. Together, our data suggest that disruption of lysosomal physiology and function are not completely responsible for impeded clearance of p62 and NBR1.

The endosomal system is a complex network heavily regulated by Rab proteins that are responsible for vesicle trafficking, fusion and maturation (*105*). These proteins are also involved in the regulation of autophagosome formation, maturation and fusion (*70, 106, 107*) and disrupted endosomal trafficking can result in unsuccessful autophagy (*41, 108*). Additionally, endosomal dysfunction has been implicated in a number of pathological sequelae that occur in the DS phenotype, like AD and diabetes (*109–111*). The observation of disrupted endosomal pathways in DS would be consistent with previous reports of enlarged early endosomes and increased endocytic uptake in Tri21 models (*71, 72, 112, 113*). Rab5 is critical for endocytosis, endosomal sorting (*74*) and it also participates in autophagosome formation and closure (*73, 114–119*). Our work regarding early endosomes reveals no difference in induction of Rab5 abundance between CTL and DS fibroblasts after SS; thus, Rab5 does not follow a pattern of accumulation like p62 that would indicate early endosome dysfunction. However, co-localization of p62 and LC3B with Rab5 after SS in DS fibroblasts would suggest fusion of autophagosomes with Rab5-positive vesicles and indicates interference of endosomal trafficking in autophagy or vice versa. This result potentially explains why localization of p62 is not restricted to a single cellular compartment, but rather appears to happen throughout the cell.

Early endosomes naturally mature into late endosomes by substitution of Rab5 with Rab7 (*75*). Based on this information and intrigued by our results regarding Rab5, we evaluated the impact of SS on late endosomes in DS fibroblasts. LAMP2 is a protein located mainly in lysosomes, but also exists on the membranes of late endosomes (*66*). Therefore, the p62 and LAMP2 co-localization reported here indicates possible involvement of late endosomes in clearance dysfunction. Similar to early endosomes, co-localization of Rab7 with p62 and LC3B demonstrates fusion of autophagosomes with late endosomes. As mentioned previously, late endosomes have the ability to fuse directly with the lysosome (*80*); thus, increased autophagosome-late endosome co-localization might increase the quantity of cargo destined for lysosomal degradation and possibly delay autophagic clearance. AL accumulation can be a direct result of this suggested mechanism. More research is needed to elucidate if fusion of autophagosomes with Rab5-positive vesicles is precluding early endosome maturation to late endosomes or if random autophagosome fusion to existing early and late endosomes occurs.

Since Rab11, a marker of recycling endosomes, interacts with late endosomes and aids in late endosome and autophagosome fusion (*82*), we examined its function in SS-mediated autophagy in DS fibroblasts. Co-localization of Rab11 with p62 and LC3B confirmed fusion of autophagosomes with recycling endosomes and indicates that the autophagosome may fuse randomly with all types of endosomal vesicles. Furthermore, Rab11 abundance in DS followed a similar pattern to p62 after SS, indicating the accumulation of recycling endosomes in DS fibroblasts after SS. Rab11 accumulation can be a mechanism of cell defense, activated due to increased abundance of autophagosome cargo in late and early endosomes. It is hypothesized that Rab11 induction is a protective mechanism of the cell to clear cargo through the plasma membrane by increasing endosomal recycling. This hypothesis is based on the concept that brains from DS human samples, a DS mouse model and DS fibroblasts exhibit increased exosome secretion as a protective mechanism against endosomal dysfunction (*120*). This argument is also supported by the fact that Rab11 appears to be regulating exosome release and overexpression of a Rab11 loss-of-function mutant abrogated exosome secretion (*121*). Current research in our lab focuses on evaluating relative kinetics of SS-mediated autophagy in DS to investigate if Rab11 accumulation is an outcome of autophagosome-endosome fusion and disruption of endosomal trafficking.

Informed by the data presented in this report, we propose a model of diminished autophagic flux due to increased fusion between autophagosomes and endosomal vesicles (Figure 8). The exact mechanism that promotes this event will be the subject of future investigation. Fusion of autophagosomes with lysosomes and slow lysosomal clearance due to increased cargo occurs as well, although further research is needed to pinpoint specific signaling machinery related to problematic degradation. This proposed model is consistent with multiple examples from the literature that confirm that co-localization of endosomal-resident proteins with p62 and LC3B can occur in other models of induced autophagy and models of neurological disorders, like Lafora disease, Sanfilippo syndrome and Amyotrophic Lateral Sclerosis (*122–125*). These reports provide corroborating evidence for impaired autophagy and general PN dysfunction in DS. General endosomal dysfunction is associated with comorbidities that appear in the DS phenotype, and evaluation of this dysfunction in DS models may provide an insight to pathological mechanisms of these comorbidities (*71, 109–112, 126, 127*). Our data also highlight the need for further mechanistic research to identify specific networks that link endosomal dysfunction with inhibited autophagic clearance and subsequent PN collapse. Likewise, it is known that an “overcrowded” cellular environment can promote protein aggregation and PN disruption (*128, 129*); therefore, it may be of interest to investigate whether dysfunctional endosomal trafficking is conducive to macromolecular overcrowding (or vice versa) that promotes protein aggregation (*130, 131*) as a mechanism of pathogenesis.

**Figure 8.**
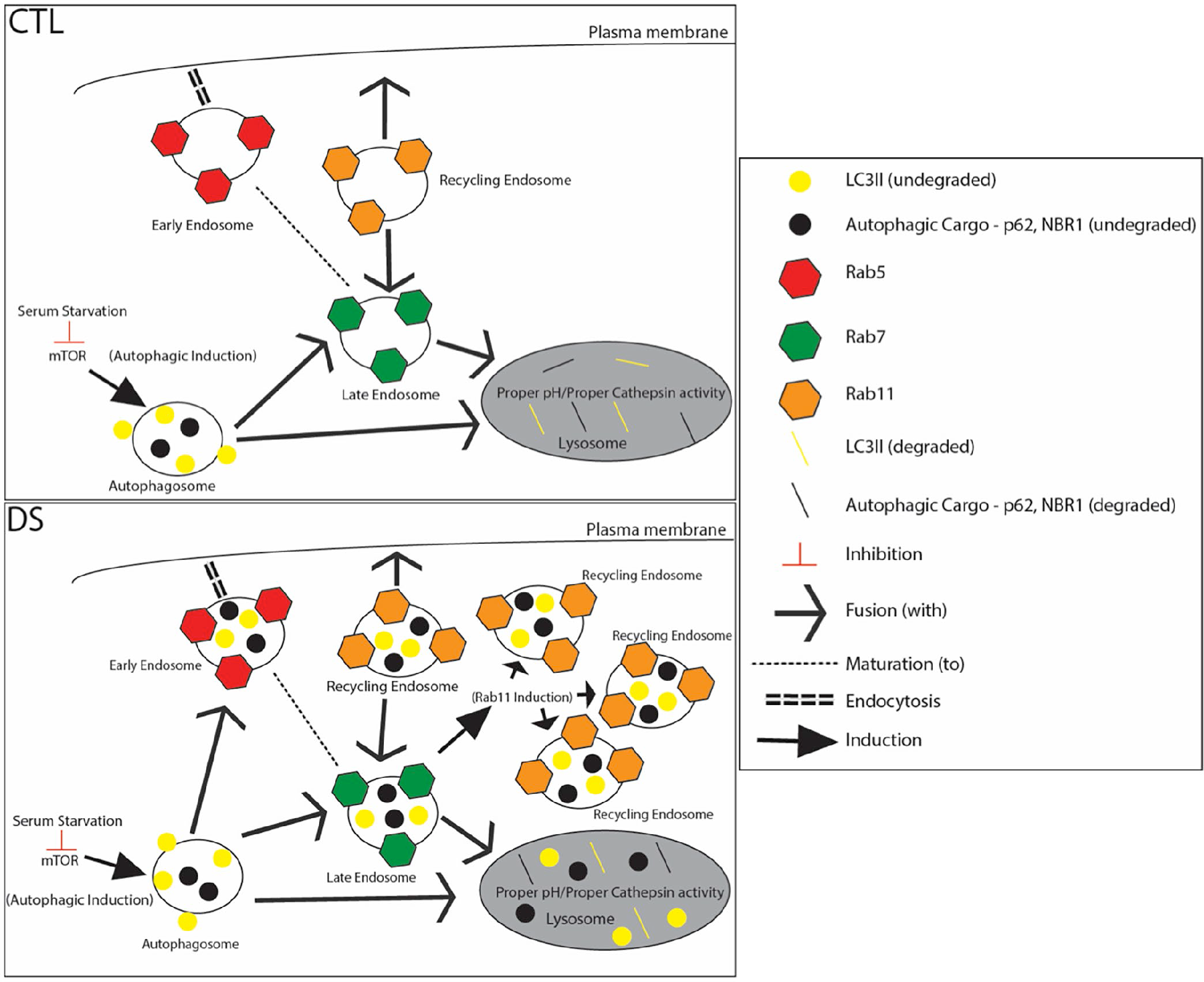
Impairment of autophagic degradation in DS fibroblasts after SS is characterized by autophagosome fusion with various endosomal vesicles and accumulation of recycling endosomes. SS inactivates mTOR and activates autophagy successfully in both CTL and DS cell lines. In the non-DS fibroblasts (CTL) the autophagosome fuses with the lysosome after SS and proper degradation occurs (degraded p62 and LC3II in the lysosome). In the DS fibroblasts (DS) the autophagosome fuses with early, late and recycling endosomes, as well as the lysosome after SS. Fusion with early and late endosomes results in an increase of the cargo destined for lysosomal degradation. Therefore, AL accumulation occurs even in the absence of lysosomal defects (Proper pH/Proper Cathepsin activity). Elevated abundance of Rab11 is also observed after SS and it is hypothesized that Rab11 induction is a cellular defense mechanism to remove cargo to the extracellular space by increasing endosomal recycling.

## Acknowledgements

The authors have no conflicts of interest to declare. This research was supported by grants from NIH/NIEHS (R01 ES027593) and the Linda Crnic Institute for Down Syndrome Research. The authors want to thank Dr. Daniel J. Klionsky (University of Michigan) and Dr. Nicholas T. Ktistakis (Babraham institute) for their valuable guidance and advice through email communication regarding experimental design. The authors also thank Dr. Rebecca McCullough for her critical review of the manuscript. Transmission electron microscopy was conducted at the Electron Microscopy Services Core Facility in the Department of Molecular Cell and Developmental Biology of University of Colorado at Boulder, with the technical assistance of facility staff.

---

**Supplemental Figure 1.**
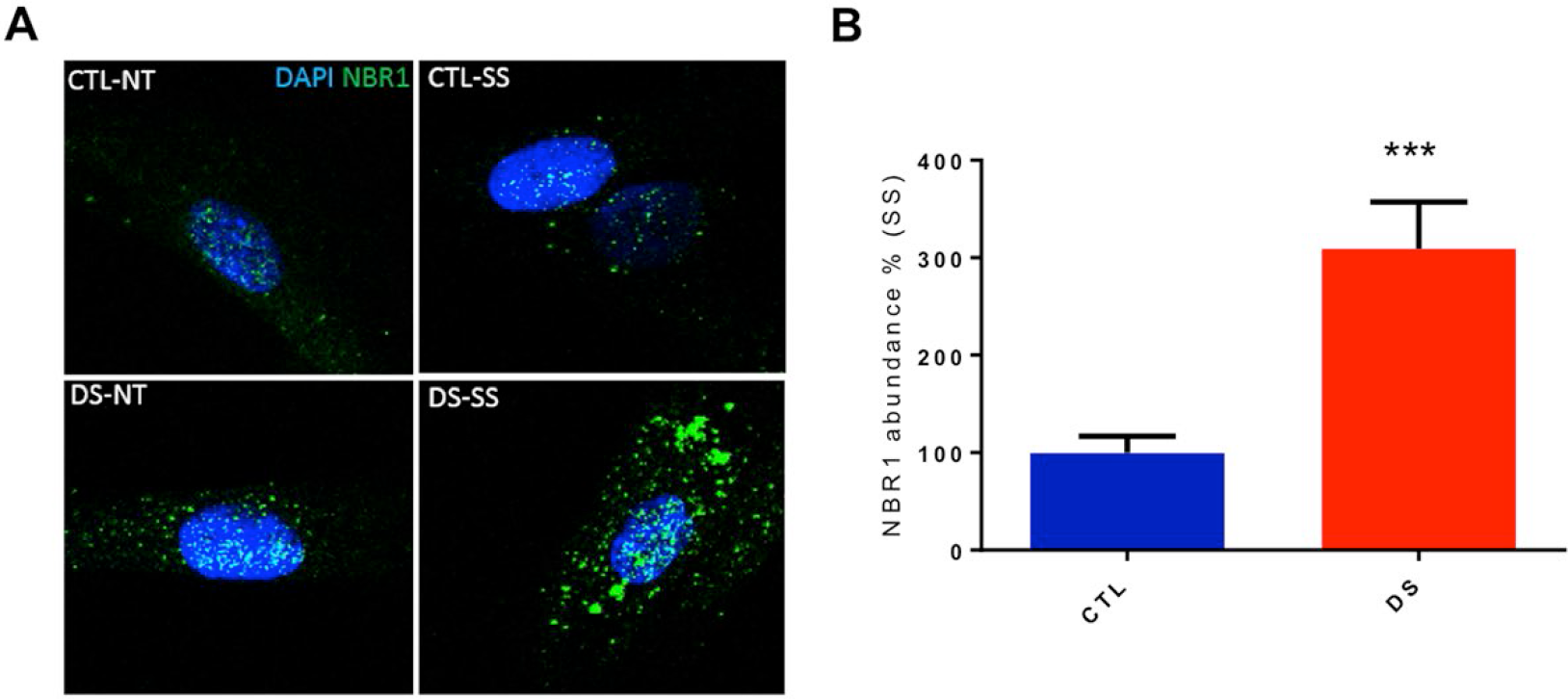
NBR1 abundance is significantly higher in DS fibroblasts after SS. (A) Immunofluorescence for NBR1 (green) and DAPI (blue) in a CTL and DS fibroblast cell line at basal levels or after SS. (B) Quantification of NBR1 (green) fluorescence intensity (area-pixels) after SS (% based on abundance of CTL SS). NT, Not treated-basal levels; SS, Serum starvation (8h).

**Supplemental Figure 2.**
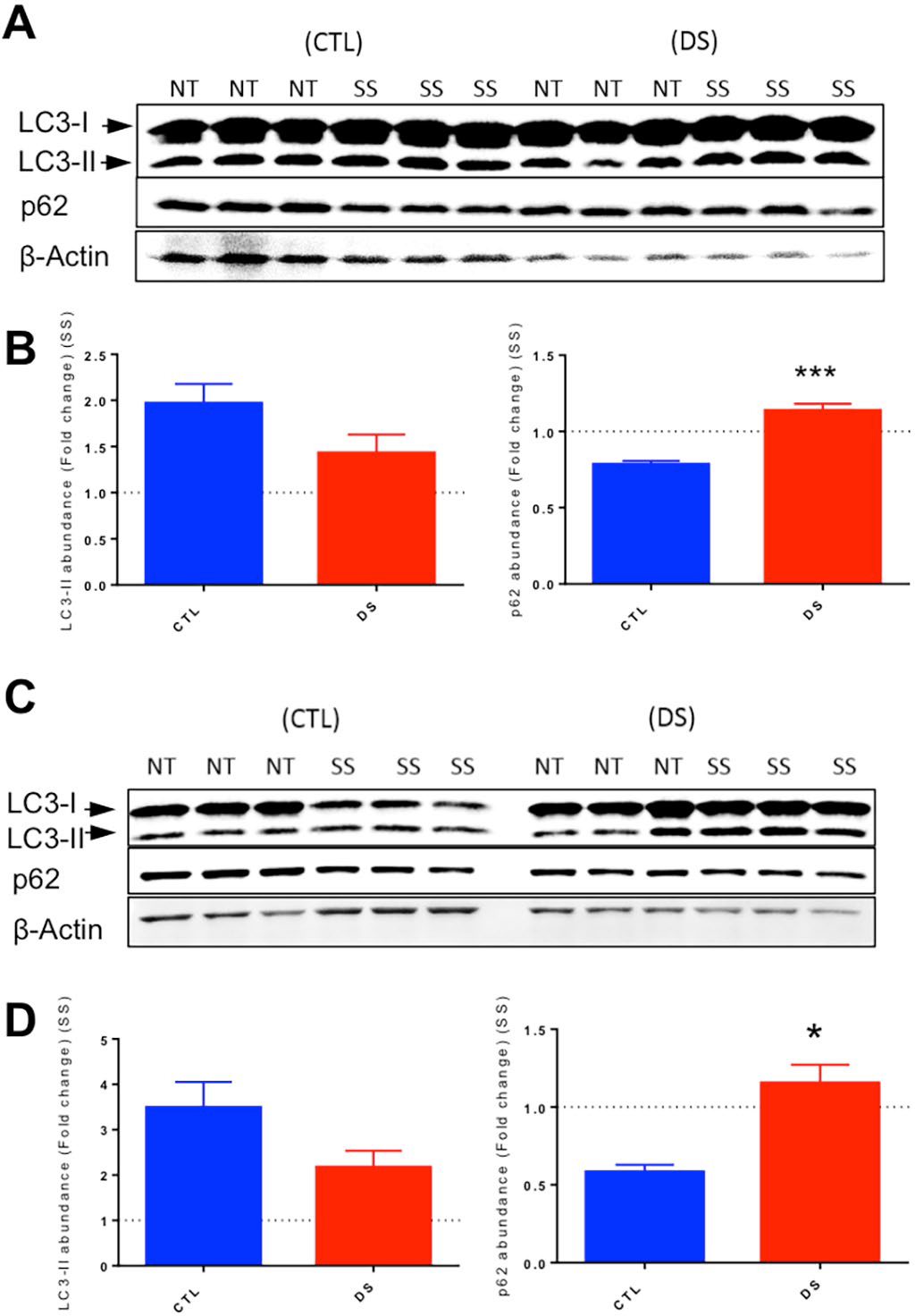
Increased fold change in p62 abundance after SS is observed in alternative DS cell models. (A) Representative blots for LC3I, LC3II, p62 and b-actin of a CTL and DS iPSC cell line at basal levels (NT) or after SS (three technical replicates per treatment and genotype are presented). (B) Quantification of fold change in abundance of p62 and LC3I-LC3II after SS. (C) Representative blots for LC3I, LC3II, p62 and b-actin of a CTL and DS NPC cell line at basal levels or after SS. (D) Quantification of fold change in abundance of p62 and LC3I-LC3II.

**Supplemental Figure 3.**
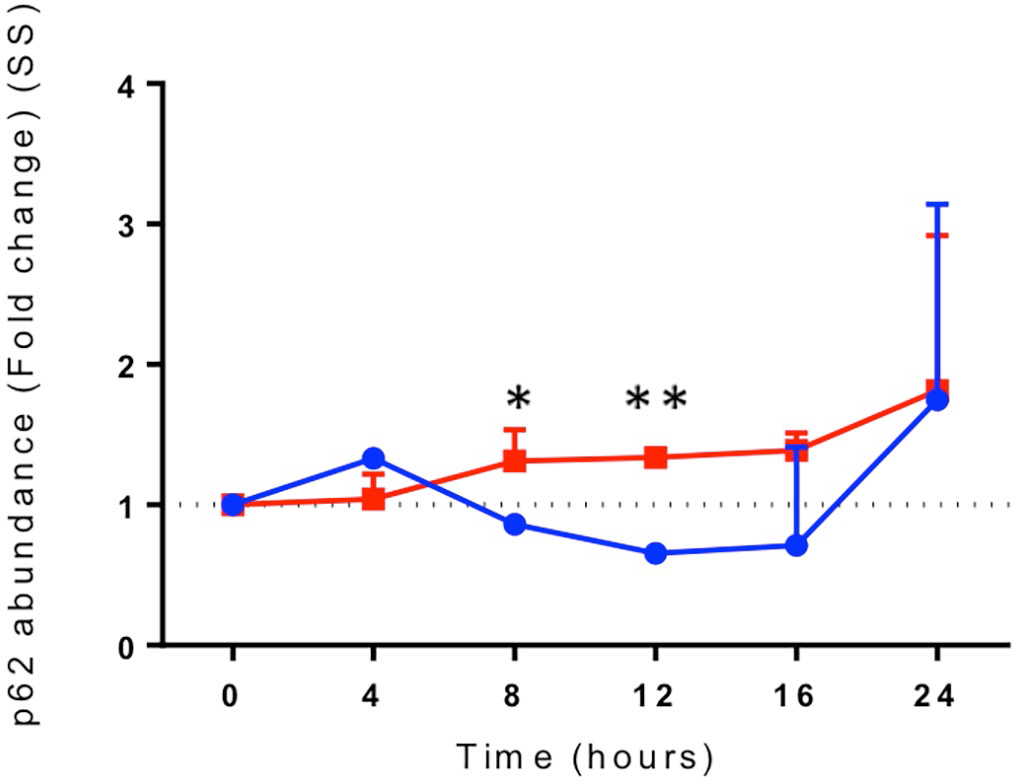
DS fibroblasts exhibit significantly increased fold change in abundance of p62 after 8h and 12h of SS compared to CTL. Western blots were conducted over a 24h period to investigate the temporal changes in p62 protein levels after serum starvation in CTL (Blue) and DS (Red) fibroblasts. Quantification of fold change in abundance of p62 at 0h, 4h, 8h, 12h, 16h or 24h of SS. Statistical analysis was performed by paired t-test analysis at each individual time point.

**Supplemental Figure 4.**
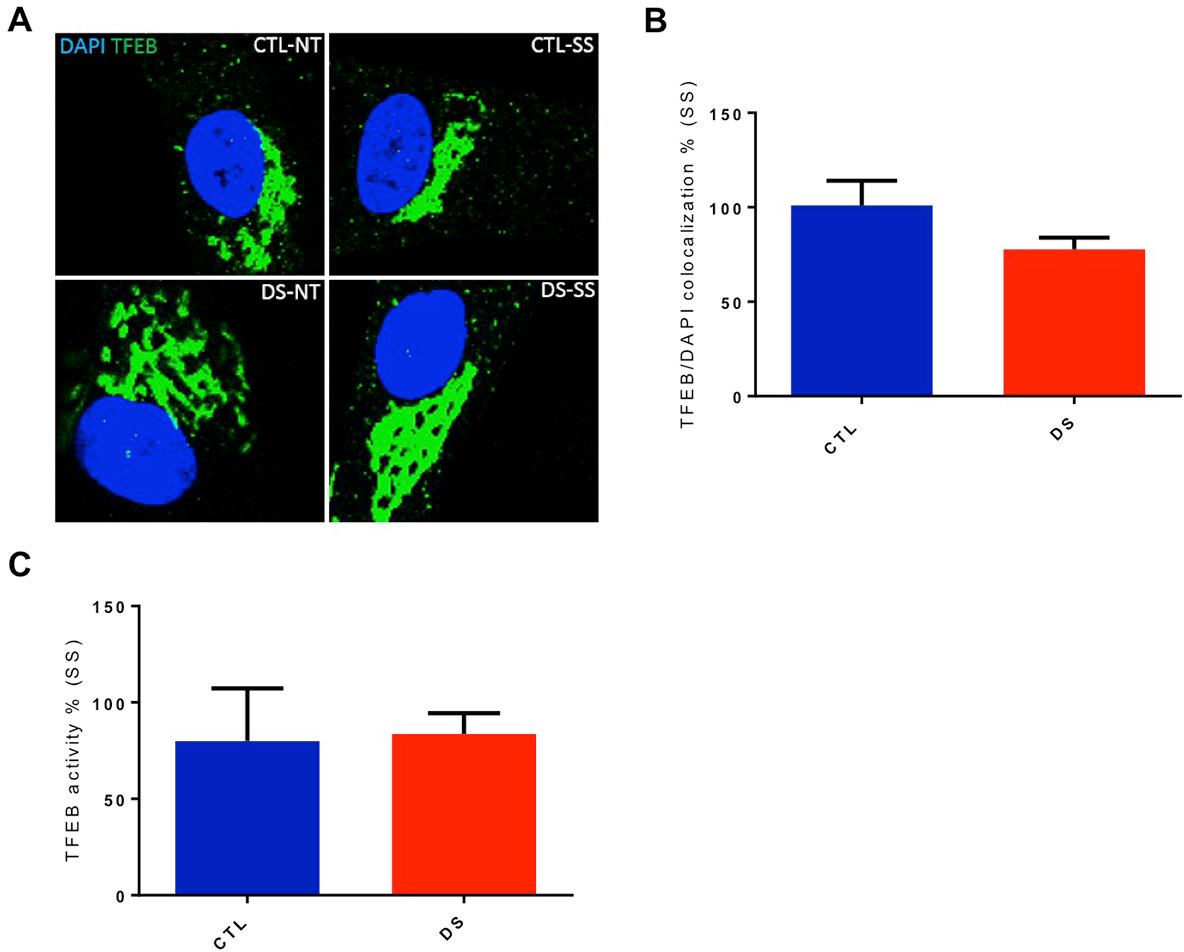
TFEB levels in the nucleus as well as TFEB DNA-binding activity are similar between CTL and DS after SS. (A) IF for TFEB (Green) and DAPI (Blue) in a CTL and DS fibroblast cell line at basal levels or after SS. (B) Quantification of TFEB/DAPI colocalization fluorescence intensity (area-pixels) after SS (% based on abundance of CTL SS). (C) Quantification of TFEB DNA-binding activity after SS (% based on activity of CTL SS). NT=Not treated SS=Serum starvation (8h)

## Abbreviations

AD: Alzheimer’s disease
AL: Autophagolysosome
CQ: Chloroquine diphosphate
CTL: Euploid control
DS: Down syndrome
HD: Huntington’s disease
IF: Immunofluorescence/Immunocytochemistry
iPSC: Induced pluripotent stem cell
mTOR: Mammalian target of Rapamycin
NPCs: Neuroprogenitor cells
NT: Basal levels-not treated
PD: Parkinson’s disease
PN: Proteostasis network
SS: serum starvation
TEM: Transmission electron microscopy
Tri8: Trisomy 8
Tri18: Trisomy 18
Tri21: Trisomy21

## References

1. F. K. Wiseman et al., A genetic cause of Alzheimer disease: mechanistic insights from Down syndrome. Nat Rev Neurosci 16, 564–574 (2015).

2. N. S. Boghossian et al., Mortality and morbidity of VLBW infants with trisomy 13 or trisomy 18. Pediatrics 133, 226–235 (2014).

3. X. Wang, T. Huang, G. Bu, H. Xu, Dysregulation of protein trafficking in neurodegeneration. Mol Neurodegener 9, 31 (2014).

4. R. J. Aitken et al., Early-onset, coexisting autoimmunity and decreased HLA-mediated susceptibility are the characteristics of diabetes in Down syndrome. Diabetes Care 36, 1181–1185 (2013).

5. H. Hasle, J. M. Friedman, J. H. Olsen, S. A. Rasmussen, Low risk of solid tumors in persons with Down syndrome. Genet Med 18, 1151–1157 (2016).

6. M. K. Mateos, D. Barbaric, S. A. Byatt, R. Sutton, G. M. Marshall, Down syndrome and leukemia: insights into leukemogenesis and translational targets. Transl Pediatr 4, 76–92 (2015).

7. C. M. Zwaan et al., Different drug sensitivity profiles of acute myeloid and lymphoblastic leukemia and normal peripheral blood mononuclear cells in children with and without Down syndrome. Blood 99, 245–251 (2002).

8. S. Aivazidis et al., The burden of trisomy 21 disrupts the proteostasis network in Down syndrome. PLoS One 12, e0176307 (2017).

9. M. Perluigi et al., Neuropathological role of PI3K/Akt/mTOR axis in Down syndrome brain. Biochimica et biophysica acta 1842, 1144–1153 (2014).

10. F. Di Domenico et al., Impairment of proteostasis network in Down syndrome prior to the development of Alzheimer’s disease neuropathology: redox proteomics analysis of human brain. Biochimica et biophysica acta 1832, 1249–1259 (2013).

11. M. Perluigi, F. Di Domenico, D. A. Buttterfield, Unraveling the complexity of neurodegeneration in brains of subjects with Down syndrome: insights from proteomics. Proteomics Clin Appl 8, 73–85 (2014).

12. C. Lanzillotta et al., Early and Selective Activation and Subsequent Alterations to the Unfolded Protein Response in Down Syndrome Mouse Models. J Alzheimers Dis 62, 347–359 (2018).

13. J. Labbadia, R. I. Morimoto, The biology of proteostasis in aging and disease. Annu Rev Biochem 84, 435–464 (2015).

14. K. Kliková et al., The Role of Heat Shock Proteins in Leukemia. Klin Onkol 29, 29–38 (2016).

15. B. Kharabi Masouleh et al., Mechanistic rationale for targeting the unfolded protein response in pre-B acute lymphoblastic leukemia. Proceedings of the National Academy of Sciences of the United States of America 111, E2219–2228 (2014).

16. J. Sun et al., Proinsulin misfolding and endoplasmic reticulum stress during the development and progression of diabetes. Molecular aspects of medicine 42, 105–118 (2015).

17. S. Jaisson, P. Gillery, Impaired proteostasis: role in the pathogenesis of diabetes mellitus. Diabetologia 57, 1517–1527 (2014).

18. Z. F. Chen et al., The double-edged effect of autophagy in pancreatic beta cells and diabetes. Autophagy 7, 12–16 (2011).

19. Y. A. Sulistio, K. Heese, The Ubiquitin-Proteasome System and Molecular Chaperone Deregulation in Alzheimer’s Disease. Mol Neurobiol 53, 905–931 (2016).

20. J. Q. Li, J. T. Yu, T. Jiang, L. Tan, Endoplasmic reticulum dysfunction in Alzheimer’s disease. Molecular neurobiology 51, 383–395 (2015).

21. A. Salminen et al., Emerging role of p62/sequestosome-1 in the pathogenesis of Alzheimer’s disease. Prog Neurobiol 96, 87–95 (2012).

22. D. Mijaljica, M. Prescott, R. J. Devenish, Microautophagy in mammalian cells: revisiting a 40-year-old conundrum. Autophagy 7, 673–682 (2011).

23. S. Kaushik, A. M. Cuervo, The coming of age of chaperone-mediated autophagy. Nat Rev Mol Cell Biol 19, 365–381 (2018).

24. C. F. Bento et al., Mammalian Autophagy: How Does It Work? Annu Rev Biochem 85, 685–713 (2016).

25. K. Dokladny, O. B. Myers, P. L. Moseley, Heat shock response and autophagy--cooperation and control. Autophagy 11, 200–213 (2015).

26. M. R. Aburto, J. M. Hurlé, I. Varela-Nieto, M. Magariños, Autophagy during vertebrate development. Cells 1, 428–448 (2012).

27. H. Li et al., Defective autophagy in osteoblasts induces endoplasmic reticulum stress and causes remarkable bone loss. Autophagy, (2018).

28. S. Di Bartolomeo, F. Nazio, F. Cecconi, The role of autophagy during development in higher eukaryotes. Traffic 11, 1280–1289 (2010).

29. G. M. Fimia, G. Kroemer, M. Piacentini, Molecular mechanisms of selective autophagy. Cell Death Differ 20, 1–2 (2013).

30. M. Pajares et al., Redox control of protein degradation. Redox Biol 6, 409–420 (2015).

31. C. Gretzmeier et al., Degradation of protein translation machinery by amino acid starvation-induced macroautophagy. Autophagy 13, 1064–1075 (2017).

32. S. Mukhopadhyay et al., Serum starvation induces anti-apoptotic cIAP1 to promote mitophagy through ubiquitination. Biochem Biophys Res Commun 479, 940–946 (2016).

33. S. T. Shibutani, T. Saitoh, H. Nowag, C. Münz, T. Yoshimori, Autophagy and autophagy-related proteins in the immune system. Nat Immunol 16, 1014–1024 (2015).

34. Y. Ma, L. Galluzzi, L. Zitvogel, G. Kroemer, Autophagy and cellular immune responses. Immunity 39, 211–227 (2013).

35. J. Lee, S. Giordano, J. Zhang, Autophagy, mitochondria and oxidative stress: cross-talk and redox signalling. Biochem J 441, 523–540 (2012).

36. A. J. Meijer, S. Lorin, E. F. Blommaart, P. Codogno, Regulation of autophagy by amino acids and MTOR-dependent signal transduction. Amino Acids 47, 2037–2063 (2015).

37. S. Sarkar, Regulation of autophagy by mTOR-dependent and mTOR-independent pathways: autophagy dysfunction in neurodegenerative diseases and therapeutic application of autophagy enhancers. Biochem Soc Trans 41, 1103–1130 (2013).

38. D. C. Rubinsztein, P. Codogno, B. Levine, Autophagy modulation as a potential therapeutic target for diverse diseases. Nat Rev Drug Discov 11, 709–730 (2012).

39. S. Sarkar, D. C. Rubinsztein, Huntington’s disease: degradation of mutant huntingtin by autophagy. FEBS J 275, 4263–4270 (2008).

40. R. A. Nixon, Amyloid precursor protein and endosomal-lysosomal dysfunction in Alzheimer’s disease: inseparable partners in a multifactorial disease. FASEB J 31, 2729–2743 (2017).

41. L. S. Whyte, A. A. Lau, K. M. Hemsley, J. J. Hopwood, T. J. Sargeant, Endo-lysosomal and autophagic dysfunction: a driving factor in Alzheimer’s disease? J Neurochem 140, 703–717 (2017).

42. M. A. Lynch-Day, K. Mao, K. Wang, M. Zhao, D. J. Klionsky, The role of autophagy in Parkinson’s disease. Cold Spring Harb Perspect Med 2, a009357 (2012).

43. A. B. Oromendia, S. E. Dodgson, A. Amon, Aneuploidy causes proteotoxic stress in yeast. Genes Dev 26, 2696–2708 (2012).

44. N. Donnelly, Z. Storchová, Aneuploidy and proteotoxic stress in cancer. Mol Cell Oncol 2, e976491 (2015).

45. A. B. Oromendia, A. Amon, Aneuploidy: implications for protein homeostasis and disease. Dis Model Mech 7, 15–20 (2014).

46. N. Dephoure et al., Quantitative proteomic analysis reveals posttranslational responses to aneuploidy in yeast. Elife 3, e03023 (2014).

47. S. Santaguida, E. Vasile, E. White, A. Amon, Aneuploidy-induced cellular stresses limit autophagic degradation. Genes Dev 29, 2010–2021 (2015).

48. Y. C. Tang, B. R. Williams, J. J. Siegel, A. Amon, Identification of aneuploidy-selective antiproliferation compounds. Cell 144, 499–512 (2011).

49. N. Donnelly, Z. Storchová, Causes and consequences of protein folding stress in aneuploid cells. Cell Cycle 14, 495–501 (2015).

50. N. Donnelly, V. Passerini, M. Dürrbaum, S. Stingele, Z. Storchová, HSF1 deficiency and impaired HSP90-dependent protein folding are hallmarks of aneuploid human cells. EMBO J 33, 2374–2387 (2014).

51. S. Stingele et al., Global analysis of genome, transcriptome and proteome reveals the response to aneuploidy in human cells. Mol Syst Biol 8, 608 (2012).

52. J. Lepenies et al., Renal TLR4 mRNA expression correlates with inflammatory marker MCP-1 and profibrotic molecule TGF-beta(1) in patients with chronic kidney disease. Nephron Clin Pract 119, c97–c104 (2011).

53. V. D. de Mello et al., Fasting serum hippuric acid is elevated after bilberry (Vaccinium myrtillus) consumption and associates with improvement of fasting glucose levels and insulin secretion in persons at high risk of developing type 2 diabetes. Molecular nutrition & food research 61, (2017).

54. L. B. Li et al., Trisomy correction in Down syndrome induced pluripotent stem cells. Cell Stem Cell 11, 615–619 (2012).

55. D. J. Orlicky, J. Monks, A. L. Stefanski, J. L. McManaman, Dynamics and molecular determinants of cytoplasmic lipid droplet clustering and dispersion. PloS one 8, e66837 (2013).

56. K. R. Parzych, D. J. Klionsky, An overview of autophagy: morphology, mechanism, and regulation. Antioxid Redox Signal 20, 460–473 (2014).

57. T. Johansen, T. Lamark, Selective autophagy mediated by autophagic adapter proteins. Autophagy 7, 279–296 (2011).

58. Y. Katsuragi, Y. Ichimura, M. Komatsu, p62/SQSTM1 functions as a signaling hub and an autophagy adaptor. FEBS J 282, 4672–4678 (2015).

59. D. J. Klionsky et al., Guidelines for the use and interpretation of assays for monitoring autophagy (3rd edition). Autophagy 12, 1–222 (2016).

60. B. E. Riley, S. E. Kaiser, R. R. Kopito, Autophagy inhibition engages Nrf2-p62 Ub-associated signaling. Autophagy 7, 338–340 (2011).

61. W. Fan et al., Keap1 facilitates p62-mediated ubiquitin aggregate clearance via autophagy. Autophagy 6, 614–621 (2010).

62. M. Komatsu et al., The selective autophagy substrate p62 activates the stress responsive transcription factor Nrf2 through inactivation of Keap1. Nat Cell Biol 12, 213–223 (2010).

63. Y. Mizunoe et al., Involvement of lysosomal dysfunction in autophagosome accumulation and early pathologies in adipose tissue of obese mice. Autophagy 13, 642–653 (2017).

64. C. Sarkar et al., Impaired autophagy flux is associated with neuronal cell death after traumatic brain injury. Autophagy 10, 2208–2222 (2014).

65. M. Redmann et al., Inhibition of autophagy with bafilomycin and chloroquine decreases mitochondrial quality and bioenergetic function in primary neurons. Redox Biol 11, 73–81 (2017).

66. Y. Endo, A. Furuta, I. Nishino, Danon disease: a phenotypic expression of LAMP-2 deficiency. Acta Neuropathol 129, 391–398 (2015).

67. E. L. Eskelinen, Y. Tanaka, P. Saftig, At the acidic edge: emerging functions for lysosomal membrane proteins. Trends Cell Biol 13, 137–145 (2003).

68. L. DeVorkin, S. M. Gorski, LysoTracker staining to aid in monitoring autophagy in Drosophila. Cold Spring Harb Protoc 2014, 951–958 (2014).

69. N. Raben, R. Puertollano, TFEB and TFE3: Linking Lysosomes to Cellular Adaptation to Stress. Annu Rev Cell Dev Biol 32, 255–278 (2016).

70. X. Ao, L. Zou, Y. Wu, Regulation of autophagy by the Rab GTPase network. Cell Death Differ 21, 348–358 (2014).

71. S. Kim et al., Evidence that the rab5 effector APPL1 mediates APP-βCTF-induced dysfunction of endosomes in Down syndrome and Alzheimer’s disease. Mol Psychiatry 21, 707–716 (2016).

72. Y. Jiang et al., Partial BACE1 reduction in a Down syndrome mouse model blocks Alzheimer-related endosomal anomalies and cholinergic neurodegeneration: role of APP-CTF. Neurobiol Aging 39, 90–98 (2016).

73. F. Zhou et al., A Rab5 GTPase module is important for autophagosome closure. PLoS Genet 13, e1007020 (2017).

74. Y. Aikawa, S. Lee, Role of Rabex-5 in the sorting of ubiquitinated cargo at an early stage in the endocytic pathway. Commun Integr Biol 6, e24463 (2013).

75. J. Huotari, A. Helenius, Endosome maturation. EMBO J 30, 3481–3500 (2011).

76. J. Rink, E. Ghigo, Y. Kalaidzidis, M. Zerial, Rab conversion as a mechanism of progression from early to late endosomes. Cell 122, 735–749 (2005).

77. T. O. Berg, M. Fengsrud, P. E. Strømhaug, T. Berg, P. O. Seglen, Isolation and characterization of rat liver amphisomes. Evidence for fusion of autophagosomes with both early and late endosomes. J Biol Chem 273, 21883–21892 (1998).

78. C. M. Fader, D. Sánchez, M. Furlán, M. I. Colombo, Induction of autophagy promotes fusion of multivesicular bodies with autophagic vacuoles in k562 cells. Traffic 9, 230–250 (2008).

79. P. A. Vanlandingham, B. P. Ceresa, Rab7 regulates late endocytic trafficking downstream of multivesicular body biogenesis and cargo sequestration. J Biol Chem 284, 12110–12124 (2009).

80. J. P. Luzio, S. R. Gray, N. A. Bright, Endosome-lysosome fusion. Biochem Soc Trans 38, 1413–1416 (2010).

81. J. P. Luzio, M. D. Parkinson, S. R. Gray, N. A. Bright, The delivery of endocytosed cargo to lysosomes. Biochem Soc Trans 37, 1019–1021 (2009).

82. Z. Szatmári et al., Rab11 facilitates cross-talk between autophagy and endosomal pathway through regulation of Hook localization. Mol Biol Cell 25, 522–531 (2014).

83. A. Longatti et al., TBC1D14 regulates autophagosome formation via Rab11- and ULK1-positive recycling endosomes. J Cell Biol 197, 659–675 (2012).

84. B. M. Frost, G. Gustafsson, R. Larsson, P. Nygren, G. Lönnerholm, Cellular cytotoxic drug sensitivity in children with acute leukemia and Down’s syndrome: an explanation to differences in clinical outcome? Leukemia 14, 943–944 (2000).

85. B. L. Bloemers, G. M. van Bleek, J. L. Kimpen, L. Bont, Distinct abnormalities in the innate immune system of children with Down syndrome. J Pediatr 156, 804–809, 809.e801-809.e805 (2010).

86. G. Ram, J. Chinen, Infections and immunodeficiency in Down syndrome. Clin Exp Immunol 164, 9–16 (2011).

87. A. Ruparelia, M. L. Pearn, W. C. Mobley, Aging and intellectual disability: insights from mouse models of Down syndrome. Dev Disabil Res Rev 18, 43–50 (2013).

88. V. Rodríguez-Sureda, Á. Vilches, O. Sánchez, L. Audí, C. Domínguez, Intracellular oxidant activity, antioxidant enzyme defense system, and cell senescence in fibroblasts with trisomy 21. Oxid Med Cell Longev 2015, 509241 (2015).

89. M. Perluigi, D. A. Butterfield, Oxidative Stress and Down Syndrome: A Route toward Alzheimer-Like Dementia. Curr Gerontol Geriatr Res 2012, 724904 (2012).

90. N. C. Firth et al., Aging related cognitive changes associated with Alzheimer’s disease in Down syndrome. Ann Clin Transl Neurol 5, 741–751 (2018).

91. E. Head, D. K. Powell, F. A. Schmitt, Metabolic and Vascular Imaging Biomarkers in Down Syndrome Provide Unique Insights Into Brain Aging and Alzheimer Disease Pathogenesis. Front Aging Neurosci 10, 191 (2018).

92. E. Bayen, K. L. Possin, Y. Chen, L. Cleret de Langavant, K. Yaffe, Prevalence of Aging, Dementia, and Multimorbidity in Older Adults With Down Syndrome. JAMA Neurol, (2018).

93. A. M. Iyer et al., mTOR Hyperactivation in down syndrome hippocampus appears early during development. J Neuropathol Exp Neurol 73, 671–683 (2014).

94. L. Rué et al., Brain region- and age-dependent dysregulation of p62 and NBR1 in a mouse model of Huntington’s disease. Neurobiol Dis 52, 219–228 (2013).

95. M. C. Micsenyi, J. Sikora, G. Stephney, K. Dobrenis, S. U. Walkley, Lysosomal membrane permeability stimulates protein aggregate formation in neurons of a lysosomal disease. J Neurosci 33, 10815–10827 (2013).

96. J. Y. Lee, Y. Nagano, J. P. Taylor, K. L. Lim, T. P. Yao, Disease-causing mutations in parkin impair mitochondrial ubiquitination, aggregation, and HDAC6-dependent mitophagy. J Cell Biol 189, 671–679 (2010).

97. K. Nakaso et al., Transcriptional activation of p62/A170/ZIP during the formation of the aggregates: possible mechanisms and the role in Lewy body formation in Parkinson’s disease. Brain Res 1012, 42–51 (2004).

98. R. M. Naylor, J. M. van Deursen, Aneuploidy in Cancer and Aging. Annu Rev Genet 50, 45–66 (2016).

99. J. W. Taub, Y. Ge, Down syndrome, drug metabolism and chromosome 21. Pediatric blood & cancer 44, 33–39 (2005).

100. D. Nižetić, J. Groet, Tumorigenesis in Down’s syndrome: big lessons from a small chromosome. Nat Rev Cancer 12, 721–732 (2012).

101. A. Cavadino, J. K. Morris, Revised estimates of the risk of fetal loss following a prenatal diagnosis of trisomy 13 or trisomy 18. Am J Med Genet A 173, 953–958 (2017).

102. E. B. Hook, D. Warburton, Turner syndrome revisited: review of new data supports the hypothesis that all viable 45,X cases are cryptic mosaics with a rescue cell line, implying an origin by mitotic loss. Hum Genet 133, 417–424 (2014).

103. X. Li et al., A molecular mechanism to regulate lysosome motility for lysosome positioning and tubulation. Nat Cell Biol 18, 404–417 (2016).

104. M. P. Nelson et al., Autophagy-lysosome pathway associated neuropathology and axonal degeneration in the brains of alpha-galactosidase A-deficient mice. Acta Neuropathol Commun 2, 20 (2014).

105. H. Stenmark, Rab GTPases as coordinators of vesicle traffic. Nat Rev Mol Cell Biol 10, 513–525 (2009).

106. Z. Szatmári, M. Sass, The autophagic roles of Rab small GTPases and their upstream regulators: a review. Autophagy 10, 1154–1166 (2014).

107. A. Kern, I. Dikic, C. Behl, The integration of autophagy and cellular trafficking pathways via RAB GAPs. Autophagy 11, 2393–2397 (2015).

108. S. A. Tooze, A. Abada, Z. Elazar, Endocytosis and autophagy: exploitation or cooperation? Cold Spring Harb Perspect Biol 6, a018358 (2014).

109. Z. Yang et al., Functional characterization of retromer in GLUT4 storage vesicle formation and adipocyte differentiation. FASEB J 30, 1037–1050 (2016).

110. R. Govers, A. C. Coster, D. E. James, Insulin increases cell surface GLUT4 levels by dose dependently discharging GLUT4 into a cell surface recycling pathway. Mol Cell Biol 24, 6456–6466 (2004).

111. S. D. Ginsberg et al., Regional selectivity of rab5 and rab7 protein upregulation in mild cognitive impairment and Alzheimer’s disease. J Alzheimers Dis 22, 631–639 (2010).

112. A. M. Cataldo et al., Down syndrome fibroblast model of Alzheimer-related endosome pathology: accelerated endocytosis promotes late endocytic defects. Am J Pathol 173, 370–384 (2008).

113. J. C. Cossec et al., Trisomy for synaptojanin1 in Down syndrome is functionally linked to the enlargement of early endosomes. Hum Mol Genet 21, 3156–3172 (2012).

114. J. P. Gorvel, P. Chavrier, M. Zerial, J. Gruenberg, rab5 controls early endosome fusion in vitro. Cell 64, 915–925 (1991).

115. A. Simonsen et al., EEA1 links PI(3)K function to Rab5 regulation of endosome fusion. Nature 394, 494–498 (1998).

116. T. Ohya et al., Reconstitution of Rab- and SNARE-dependent membrane fusion by synthetic endosomes. Nature 459, 1091–1097 (2009).

117. G. G. Tall, M. A. Barbieri, P. D. Stahl, B. F. Horazdovsky, Ras-activated endocytosis is mediated by the Rab5 guanine nucleotide exchange activity of RIN1. Dev Cell 1, 73–82 (2001).

118. B. Ravikumar, S. Imarisio, S. Sarkar, C. J. O’Kane, D. C. Rubinsztein, Rab5 modulates aggregation and toxicity of mutant huntingtin through macroautophagy in cell and fly models of Huntington disease. J Cell Sci 121, 1649–1660 (2008).

119. C. E. Chua, B. Q. Gan, B. L. Tang, Involvement of members of the Rab family and related small GTPases in autophagosome formation and maturation. Cell Mol Life Sci 68, 3349–3358 (2011).

120. S. A. Gauthier et al., Enhanced exosome secretion in Down syndrome brain - a protective mechanism to alleviate neuronal endosomal abnormalities. Acta Neuropathol Commun 5, 65 (2017).

121. A. Savina, M. Vidal, M. I. Colombo, The exosome pathway in K562 cells is regulated by Rab11. J Cell Sci 115, 2505–2515 (2002).

122. M. Osaka, D. Ito, T. Yagi, Y. Nihei, N. Suzuki, Evidence of a link between ubiquilin 2 and optineurin in amyotrophic lateral sclerosis. Hum Mol Genet 24, 1617–1629 (2015).

123. R. Puri, T. Suzuki, K. Yamakawa, S. Ganesh, Dysfunctions in endosomal-lysosomal and autophagy pathways underlie neuropathology in a mouse model for Lafora disease. Hum Mol Genet 21, 175–184 (2012).

124. H. Beard et al., Axonal dystrophy in the brain of mice with Sanfilippo syndrome. Exp Neurol 295, 243–255 (2017).

125. X. Chen et al., Autophagy induced by calcium phosphate precipitates targets damaged endosomes. J Biol Chem 289, 11162–11174 (2014).

126. V. Udayar et al., A paired RNAi and RabGAP overexpression screen identifies Rab11 as a regulator of β-amyloid production. Cell Rep 5, 1536–1551 (2013).

127. M. Uhlig, W. Passlack, J. Eckel, Functional role of Rab11 in GLUT4 trafficking in cardiomyocytes. Mol Cell Endocrinol 235, 1–9 (2005).

128. Z. Zhou et al., Crowded cell-like environment accelerates the nucleation step of amyloidogenic protein misfolding. J Biol Chem 284, 30148–30158 (2009).

129. M. Bokvist, G. Gröbner, Misfolding of amyloidogenic proteins at membrane surfaces: the impact of macromolecular crowding. J Am Chem Soc 129, 14848–14849 (2007).

130. A. L. Zajac, Y. E. Goldman, E. L. Holzbaur, E. M. Ostap, Local cytoskeletal and organelle interactions impact molecular-motor-driven early endosomal trafficking. Curr Biol 23, 1173–1180 (2013).

131. P. Sood et al., Cargo crowding at actin-rich regions along axons causes local traffic jams. Traffic 19, 166–181 (2018).

